# A fossil fish assemblage from the middle Miocene of the Cocinetas Basin, northern Colombia

**DOI:** 10.1101/2021.04.19.440491

**Authors:** Gustavo A. Ballen, Carlos Jaramillo, Fernando C. P. Dagosta, Mario C. C. De Pinna

## Abstract

Freshwater fossil fish faunas have been long used to infer past drainage connections, as they are bounded by physical freshwater barriers. Here we study a middle Miocene (15.0-–15.5 Ma) freshwater fish fossil fauna (Makaraipao) from the Castilletes Formation in northern Colombia, nowadays west of the Andes. We record the presence of lungfishes (*Lepidosiren*), pacus (*Mylossoma* and *Piaractus*), armored catfishes (Callichthyidae), and red-tail catfishes (*Phractocephalus*). Extant members of all those groups (except the Callichthyidae, due to lack of taxonomic resolution) are found in Amazonian faunas east of the Andes and are absent from faunas west of the Andes, indicating that the riverine systems of the Guajira Peninsula were connected to Amazonia during the middle Miocene. The similarity of La Venta (west of the Andes) and Rio Acre (east of the Andes) fish faunas during the late Miocene further indicates that the northern Andean uplift was not a complete barrier at least until *∼* 11 Myr ago. However, there is a continental-wide structuring of the Miocene fish faunas that is also found in the extant faunas, suggesting that other factors such as ecological conditions, in addition to the uplift of the Andes, have shaped the biogeographic evolution of South American fish faunas.

## INTRODUCTION

Freshwater fishes are the richest vertebrate continental component of the Neotropics, with *∼* 7,000 species (Albert & Reis 2011). They are highly diverse, ranging from brackish waters to elevations above 3000 m in the Andes (Schaefer 2011). Despite this enormous diversity, freshwater fishes are not as prominent in the fossil record of South America as mammals or crocodylians, and they are often recorded from bone fragments of limited diagnostic value (Lundberg *et al*. 2010). The paucity of wide-ranging comparative morphological analyses focused on diagnostic characters preserved in fossil fish specimens limits the identification to only coarse taxonomic levels such as order or family. Such drawbacks limit potential use of Neotropical freshwater fossil fishes in paleoecological, systematic, biostratigraphic, and biogeographic studies.

Freshwater fishes have provided evidence of ancient drainage connections between river systems east and west of the Andes, as the dispersal potential of freshwater fishes across mountain ranges is limited for lowland taxa, thus helping to constrain the evolution of the Andean orogeny (Lundberg 1997; Lundberg *et al*. 2010). For example, extant freshwater fish taxa currently restricted to cis-Andean drainages (e.g.,Orinoco-Amazon) are found in Miocene trans-Andean sites (i.e., west of modern Andes range as La Venta, Castilletes and Urumaco), indicating ancient hydrological connectivity that is severed nowadays (Diaz de Gamero 1996; Gregory-Wodzicki 2000; Montes *et al*. 2021).

A number of studies have improved our understanding of the geology and paleontology of the Cocinetas sedimentary basin in the Guajira Peninsula of northern Colombia (Moreno *et al*. 2015). Several vertebrate groups are represented in these sediments, both marine and continental (Aguilera *et al*. 2013*a*; Forasiepi *et al*. 2014; Cadena & Jaramillo 2015*a,b*; Amson *et al*. 2016; Moreno-Bernal *et al*. 2016; Suarez *et al*. 2016; Aguilera *et al*. 2017; Carrillo-Briceño *et al*. 2019); in contrast, freshwater fishes remains largely unreported in these assemblages (but see Aguilera *et al*. 2013*b*).

Herein, we report the freshwater fossil fishes of the Miocene Castilletes Formation in the Makaraipao locality, northern Colombia, and their relevant anatomical characters. We compare the fossil assemblage with South American fish faunas as well as with the extant continental fish diversity in order to understand the historical component of similarity patterns in time and space.

## MATERIALS AND METHODS

### Geological setting

The Cocinetas sedimentary basin preserves a succession of continental to shallow marine strata on the northern margin of the South American Plate. Moreno *et al*. (2015) redefined the stratigraphy of the basin and showed that the Castilletes Formation is composed of sediments of Miocene age from mostly shallow marine environments. Its lower boundary is concordant with the Jimol Formation whereas its upper boundary is set by an unconformity with the Ware Formation. The Castilletes Formation has been dated as lower to middle Miocene using Sr geochronology (Hendy *et al*. 2015).

Makaraipao (locality STRI 390093, 11°54’32.0“N 71°20’24.4”W) is a moderate plateau lying to the west of the Tucacas bay, municipality of Uribia, Guajira department, northern Colombia (Figure 1a-c). The locality is recorded in the stratigraphic section called “Long section” (Moreno *et al*. 2015). The locality lies about 127 stratigraphic meters above the base of the section (Figure 1d) and *∼* 279 m above the base of the Castilletes formation (Suarez *et al*. 2016). Specimens were collected from a sandstone and conglomeratic sandstone level. The Sr isotopic data constrains the Makaraipao locality to a 15.0–-15.5 Ma age (Carrillo-Briceño *et al*. 2019).

**FIG. 1:**
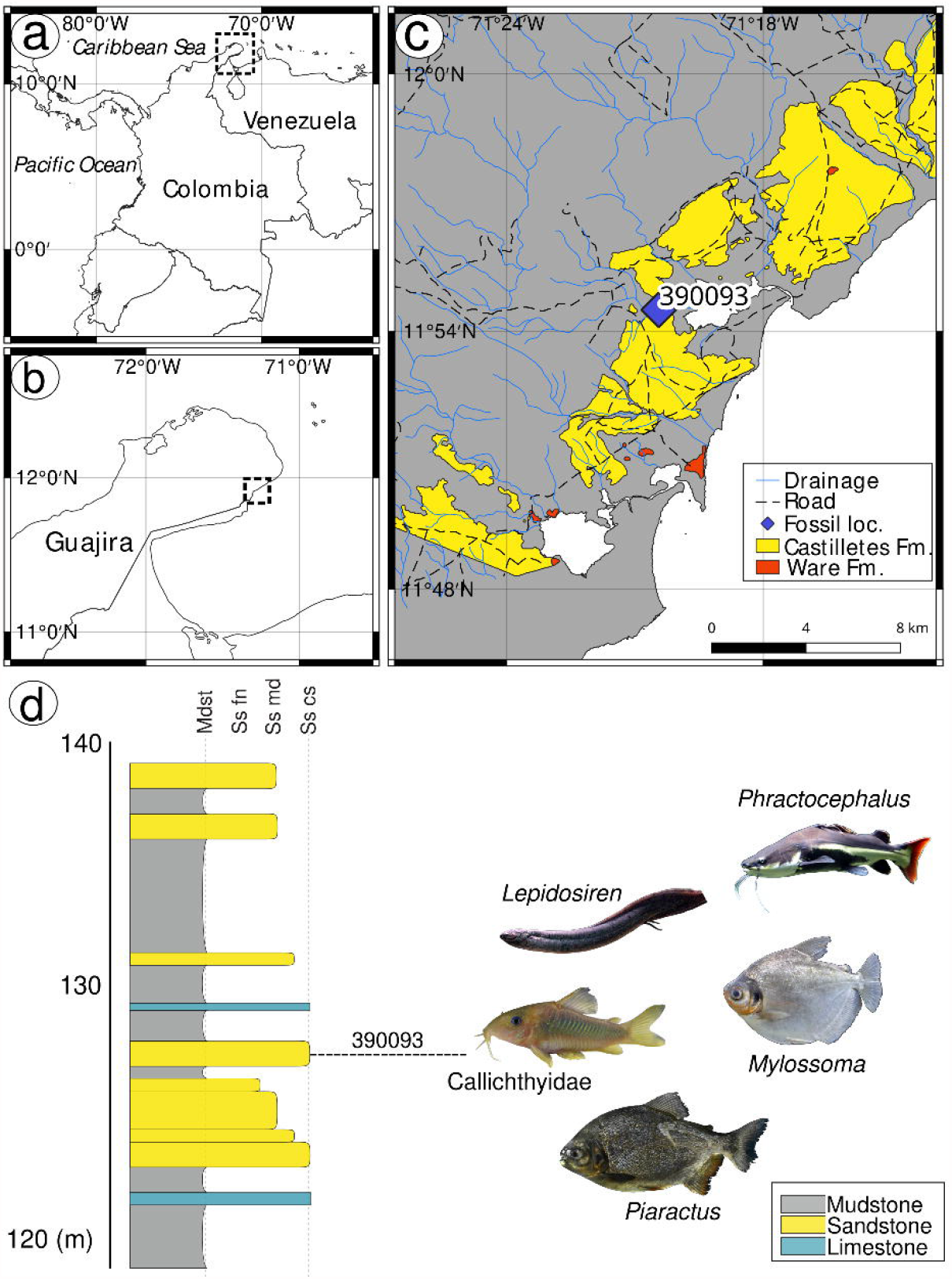
a-c) Geographic context of the locality Makaraipao in the Cocinetas Basin, Guajira Peninsula, Colombia. d) Stratigraphic column of the Long Section (loc. 170514) modified from Moreno *et al*. (2015), vertical scale in meters. Extant representatives of the taxa recorded in the fossil locality are shown to the right of the locality number. Photographs of extant representatives by Daiju Azuma (*Lepidosiren*), Carl Clifford (*Phractocephalus*), Nadia Milani (*Mylossoma*), Bruno S. Barros (*Piaractus*), and Karsten Schönherr (Callichthyidae).

### Abbreviations

Institutional abbreviations are: Academy of Natural Sciences of Drexel University, Philadelphia, US (ANSP), Instituto Alexander von Humboldt, Villa de Leyva, Colombia (IAvH), Museu de Zoologia da Universidade de São Paulo, São Paulo, Brazil (MZUSP), Mapuka Museum of Universidad del Norte, Barranquilla, Colombia (MUN). Premaxilla and dentary are abbreviated PM and D respectively. Approximately-unbiased and bootstrap nodal support values are termed AU and BT respectively.

### Anatomical terminology

Descriptive nomenclature of serrasalmid dentition follows Cione *et al*. (2009) with modifications where needed (Figure 2). Lepidosirenid anatomical terminology follows Criswell (2015). Terminology of the Siluriform appendicular follows Ballen & Pinna (2021).

**FIG. 2:**
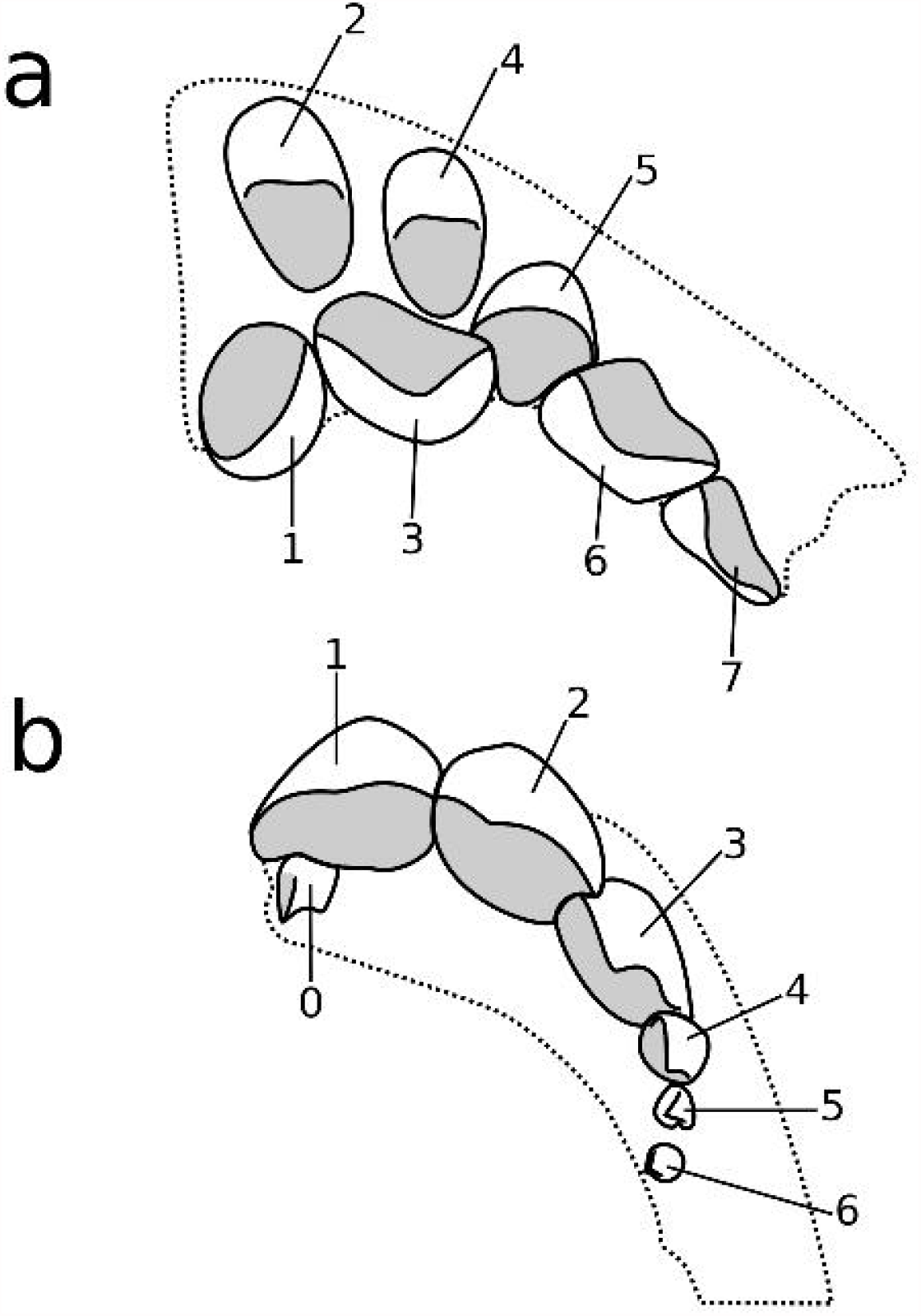
Schematic representation of a Serrasalmid Pacu dentition based on *Myloplus lucienae* (Andrade *et al*. 2016). (a) Premaxilla with teeth numbered following Cione *et al*. (2009); (b) dentary numbering as herein proposed with the digit zero to the symphysial dentary tooth. Gray shade represents occlusal molariform surfaces for teeth where such feature is present. Cutting edge shape and position represented for each tooth.

### Data analysis

Faunal composition for a similarity analysis was compiled from literature data and direct observations for the fossil assemblages Castillo, La Venta, Makaraipao, Castilletes marine, Loyola-Mangan, Rio Acre, Solimões-Pebas, Urumaco, Utuquina, Pirabas, Cantaure, Ituzaingo, Rio Yuca, Fitzcarrald, and Contamana (Supplementary Table S1) (Cione *et al*. 2000, 2009; Lundberg *et al*. 2010; Bogan *et al*. 2012; Aguilera *et al*. 2013*a,b*; Cione & Azpelicueta 2013; Tejada-Lara *et al*. 2015; Antoine *et al*. 2016; Azpelicueta & Cione 2016; Ballen & Moreno-Bernal 2019). We compiled an initial set of Miocene fossil fish fauna and later limited the analysis to those faunas that had a sampling size equal to or larger than the one herein described, in order to avoid uncertainty due to low sample size. Therefore, the faunas used in the analysis were reduced to La Venta (12.0 Ma), Makaraipao (15.0 Ma), Rio Acre (7.9 Ma), Urumaco (8.0 Ma), Ituzaingo (7.5 Ma), Fitzcarrald (12.8 Ma), and Contamanta (11.0 Ma). An analysis with the full dataset provided a similar pattern as described here (Supplementary section S1). Modern faunal similarity was measured using a continental-wide database of curated, voucher-based presence/absence records of fish species in each of the hydrographic regions of South America (Dagosta & Pinna 2019). Faunal similarity and resampling nodal support values were measured using the binary method with average distance as implemented in the pvclust package v.2.2-0 (Suzuki *et al*. 2019) in R v.3.4.4 R Core Development Team (2018). We chose the binary method because we have presence/absence instead of abundance data and because of the asymmetry with respect to absences. Resampling nodal support measures AU and BT were calculated with the pvclust::pvclust function; although both measures are reported, AU are more reliable than BT as a measure of relative support in hierarchical clustering (Suzuki *et al*. 2019). The same method of clustering was applied to both the Miocene and modern datasets. Additional correlations between similarity, geologic age, and geographic proximity were calculated using the packages vegan and sf (Pebesma 2018; Oksanen *et al*. 2019) (Supplementary Section S3).

Extant occurrences were downloaded from the SpeciesLink and GBIF databases (DOI links https://doi.org/10.15468/dl.6glxkb, https://doi.org/10.15468/dl.d33vwn, https://doi.org/10.15468/dl.67lq6f, and https://doi.org/10.15468/dl.9aryay) and specific data cleaning procedures carried out (Supplementary section S2). Mapping was carried out in QGIS v.3.4.12 (QGIS Development Team 2019). Image edition and processing was carried out in GNU image manipulation program (GIMP) and Inkscape. The raw data and scripts are available both in zenodo (doi:XXXXXXXXXXXX) and https://github.com/gaballench/makaraipao.

## RESULTS

> ***Systematic paleontology***
>
> Dipnoi
>
> Order Lepidosireniformes
>
> Genus *Lepidosiren*
>
> *Lepidosiren* sp. (Figures 3g-i)

**FIG. 3:**
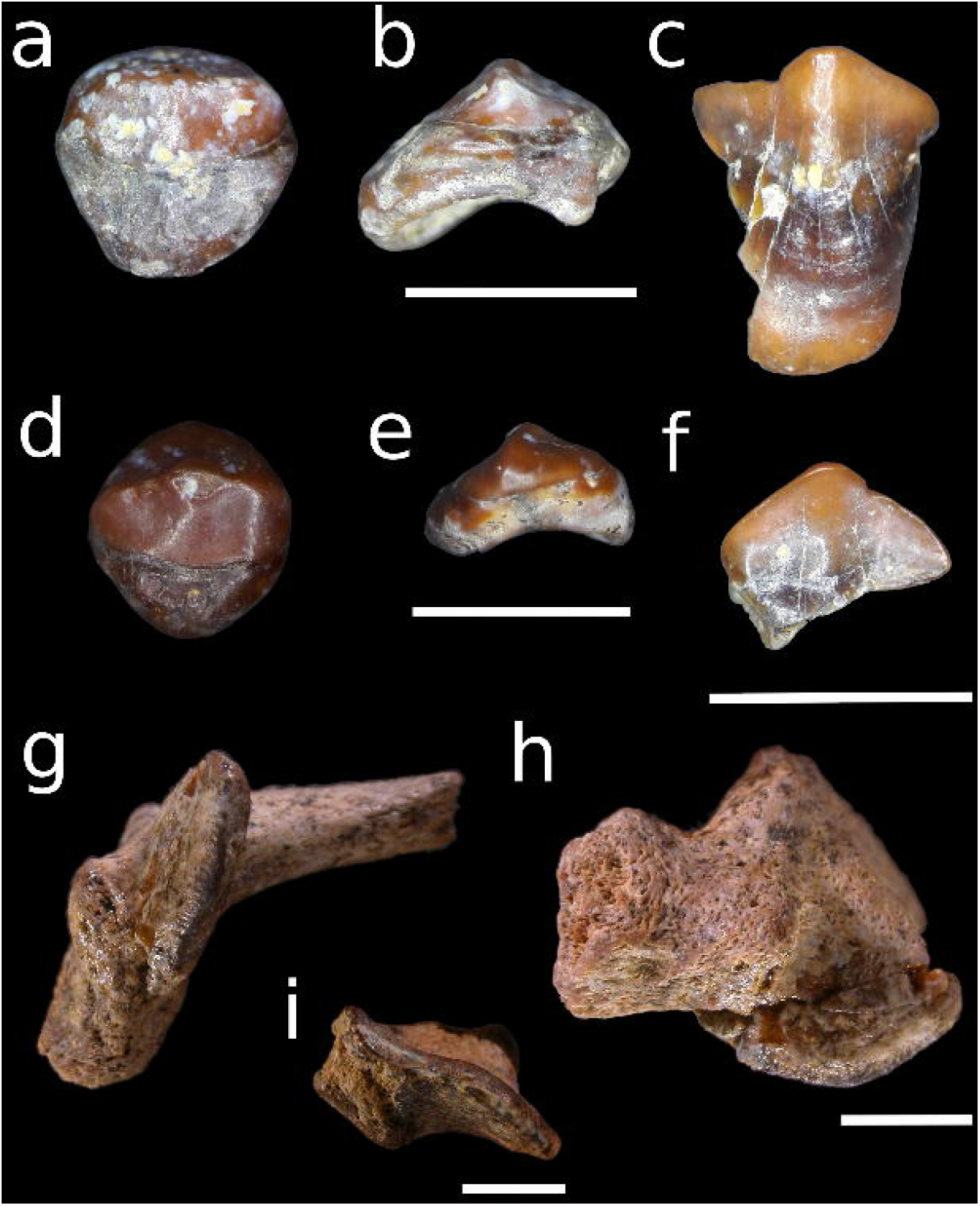
Characiforms and lepidosireniforms from the middle Castilletes Formation in the locality Makaraipao. a-b,d-e) *Piaractus* aff. *brachypomus* MUN 37664, D4-6 in occlusal (a,d) and commissural (b,e) views. c,f) *Mylossoma* sp. MUN 34502 (c,f), D2 in labial (c) and occlusal (f) views. g-i) *Lepidosiren* sp. MUN 37667 (g-h) and MUN 37693 (i) in occlusal (g,i), and labial symphysial (h) views. Scale bars equal 5 mm in all paired view.

### Material examined

MUN 37667, partial left pterygoid plate with part of middle and posterior pterygoid ridges, posterior process and base of ascending process preserved; MUN 37693, partial pterygoid plate with part of the middle pterygoid ridge and all of the posterior ridge.

### Description

Pterygoid tooth plates with middle and posterior ridges in both specimens, also with osseous support in MUN 37667. Posterior pterygoid ridge roughly sigmoid in both specimens, projecting laterally from body of pterygoid bone. Preserved crown with median portion of middle pterygoid ridge approaching contralateral ridge. Angle between ridges *∼* 30°, angle between posterior process of pterygoid and enamel-bearing axis of pterygoid *∼* 130°.

### Remarks

The pterygoid and prearticular tooth plates in Dipnoi are the functional analogues of the premaxilla and dentary (and sometimes also the maxillary and palatal teeth), respectively, in bony fishes. Criswell (2015) corroborated the South American genus *Lepidosiren* as sister group to the African *Protopterus*. The jaws of the two genera can be distinguished by the different relative proportion of the posteriormost two pterygoid ridges. The second pterygoid ridge is about half the length of the first and posteriormost ones in *Lepidosiren*, whereas the second ridge is shorter than half the length of the posteriormost one in *Protopterus*. Pterygoid tooth plates of the Lepidosirenidae are distinguished from prearticular ones by their reduced amount of enameloid, a discontinuity between the ventral outline of the pterygoid ramus and the base of the enameloid in lateral view, and the presence of an ascending process of the pterygoid (vs. ventral surface smooth and straight in lateral view) (Criswell 2015). Also, the contralateral middle ridges of the prearticular do not meet at the midline whereas they do in the pterygoid.

The extant lungfish *Lepidosiren paradoxa* is currently restricted to lentic systems, swamps, *várzeas*, and lagoons in the Amazon, Paraná, and Paraguay basins as well as in the Guyanas (Almeida-Val *et al*. 2011; Figure 4a). This taxon suggests that the fine-grained conditions recorded in levels adjacent to the sandy-conglomeratic lithology where the specimens were collected represent a peripheral lentic system.

**FIG. 4:**
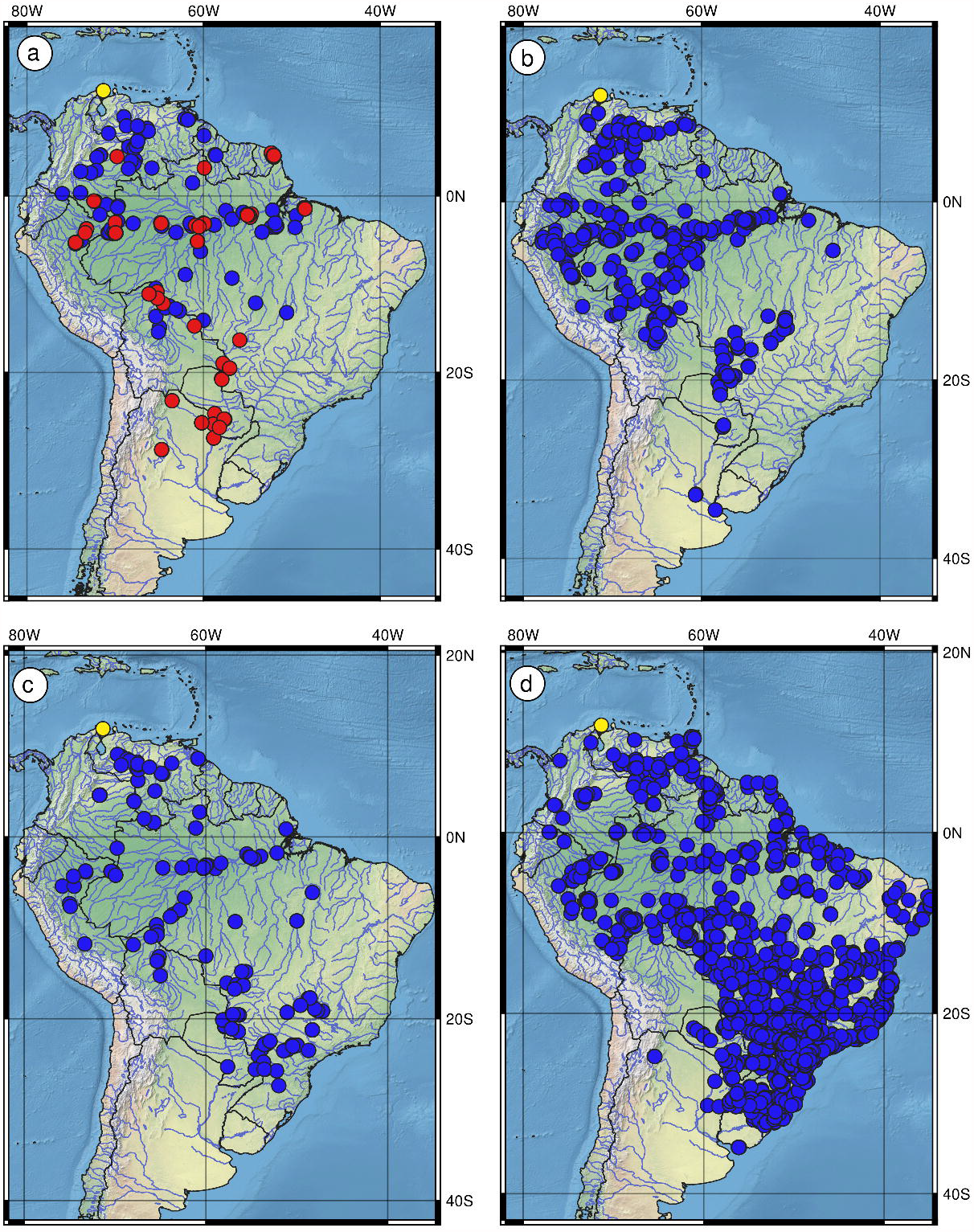
Recent distributions of fossil taxa. a) *Lepidosiren* (red) and *Phractocephalus* (blue); b) *Mylossoma*; c) *Piaractus*; d) Callichthyidae. Yellow spots in all maps represent the fossil locality Makaraipao in northern Colombia.

*Lepidosiren megalos* described by Silva Santos (1987) from the Brazilian Acre fauna, was based on cranial remains and isolated tooth plates. Lundberg *et al*. (2010) had doubts about being a different species compared to the extant *L. paradoxa* as their main difference is body size. Silva Santos (1987) also indicates that the coronoid process is shallower in *L. megalos* compared to *L. paradoxa*, but, thee difference is not evident in thepublished illustrations (Silva Santos 1987figs. 1 and 6 in plates I and II respectively). Further analyses are needed to assess if they are indeed different. The type material of *L. megalos* was allegedly housed in the paleontology collection of MZUSP but it has not been located yet (A. Carvalho & H. Britski, pers. comm., 2019). Therefore, given the lack of any published or observable distinctive characters, *Lepidosiren megalos* is herein considered a junior synonym of *Lepidosiren paradoxa*, as suggested by Lundberg *et al*. (2010), an opinion never followed in subsequent literature (Agnolin 2010; López-Fernández & Albert 2011; Alves *et al*. 2013).

*Lepidosiren* has been found in the Honda group in Colombia, the Solimões Formation in Brazil, and the Pebas Formation in both the Contamana and Fitzcarrald areas in Peru (Tejada-Lara *et al*. 2015; Antoine *et al*. 2016). Gayet *et al*. (2001) report remains of *Lepidosiren* cf. *paradoxa* from the El Molino Formation in localities of Pajcha Pata and Vila Vila in Bolivia of Danian age (early Paleocene following Gelfo *et al*. 2009; formerly thought to be Maastrichtian). These remains are the oldest record of *Lepidosiren* in South America, but their species-level identity remains elusive. A proper restudy is necessary to determine whether they are conspecific with *L. paradoxa*, however, a species timespan of *∼* 63 Myr seems unlikely.

The fossil record of the genus is consistent with the habitat of extant *Lepidosiren* including lotic waters and floodplains of the Amazon (Almeida-Val *et al*. 2011). As fossils they have been, found in the fine-grained facies of the Honda group (Ballen & Moreno-Bernal 2019).

> Division Ostariophysi
>
> Order Characiformes
>
> Family Serrasalmidae
>
> Genus *Mylossoma*
>
> *Mylossoma* sp. (Figures 3c,f)

### Material examined

MUN 34502, an isolated tooth.

### Description

Molariform D2 tooth with asymmetric crown and flat occlusal surface. Posterior margin smoothly angular, lacking strong concavities. Anterior surface with longitudinal sulcus extending from cutting edge to crown base. Crown unicuspid, with occlusal surface flat to slightly concave.

### Remarks

Multicuspid, cutting to incisiform teeth are a well-known feature of carnivore, lepidophagous, and omnivore serrasalmids (Mirande 2010; *Serrasalmus, Pygocentrus, Pristobrycon, Pygopristis*, and *Catoprion*; Kolmann *et al*. 2018). Teeth in *Megapiranha* are still reminiscent of the generalized, multicuspid, incisiform condition found in carnivore, lepidophagous, and omnivore genera (Cione *et al*. 2009). Teeth in *Acnodon, Mylesinus, Ossubtus*, and *Tometes* are multicuspid and incisiform. Contrastingly, the genera *Colossoma, Metynnis, Myleus, Mylossoma, Myloplus, Piaractus*, and *Utiaritichthys*, have molariform dentary teeth. This latter morphology is the same as that in the fossil specimen reported herein. Among recent representatives of those genera (Figure 4), only cis-Andean species of *Mylossoma* (except the trans-Andean *M. acanthogaster*) show the vertical lingual sulcus diagnostic for that cluster of species, thus permitting identification of the teeth as belonging to that genus.

The Serrasalmidae is the family with the most abundant fossil record in the Characiformes, although most of the material consists of isolated teeth that are often indeterminate beyond family level. The trophic ecology of serrasalmids is diverse, reflected in an equally diverse tooth morphology. Their dental variation is both serial (i.e., along a given tooth series in the same species) and taxonomic (among different genera and species). The greatest difficulty in using dental morphology as a taxonomic character is the lack of comparative information on extant representatives of the family. Assessing the variation at different taxonomic levels in extant taxa is therefore necessary for taxonomic identification of isolated fossil remains. The comparative study of dentary and premaxillary teeth in Recent serrasalmids that is presented here thus provides a framework for the identification of isolated fossil teeth in this fish group.

Dahdul (2004) attempted to identify fossil serrasalmid teeth at the genus level. She reports the genus *Mylossoma* from the Castillo formation in Venezuela, based on comparisons with *Colossoma* and *Piaractus* among serrasalmids, but not with other taxa with molariform teeth such as *Metynnis, Myleus*, or *Myloplus*. The teeth illustrated by Dahdul (her pl. 1) show two concavities in occlusal view, one labiolingual, associated with the lateral cusplet, and one extensive mesiolingual, associated with the tooth D0. Those characteristics are present only in *Mylossoma* among serrasalmids. Other genera have only a mesiolingual concavity (*Acnodon, Colossoma, Metynnis, Myleus, Myloplus, Piaractus*, and *Utiaritichthys*), two lingual concavities (*Myleus*), or lack concavities altogether on D1 in occlusal view (*Mylesinus* and *Tometes*). Our data thus confirm that the material from the Castillo Formation reported by Dahdul (2004) is indeed *Mylossoma* sp.

Gayet *et al*. (2001:52, fig. 7c-e) reported isolated serrasalmid teeth from the Paleocene locality of Pajcha Pata in Bolivia. Both teeth of morphotype 2 in Gayet *et al*. (2001, ig. 7d–e) are from the premaxilla, based on the presence of two cutting edges, one on the labial face and one on the lingual face. The tooth illustrated in their figure 7e can be further identified as PM3, PM6, or PM7, due to the presence of three cusps on the lingual cutting edge and at least one on the labial cutting edge.

This combination of characteristics is present only in species of *Acnodon* and *Metynnis*. Furthermore, the teeth have high labial and lingual cutting edges, a trait typical of *Acnodon* (Jégu & Santos 1990. figs. 9-10). We conclude therefore that the serrasalmid teeth from the Paleocene of Pajcha Pata reported by Gayet *et al*. (2001:52, fig. 7c-e) belong to *Acnodon*.

Some published works illustrate fossil teeth with a lingual projection in occlusal view that is bounded by two concave margins (e.g., (Rubilar 1994; Monsch 1998, plate iii fig. 13; Figure 4c). This morphology is found in PM5 across several genera where adjacent teeth compress the lingual margin thus creating two lingual concavities and examples include *Colossoma, Myloplus, Metynnis, Myleus, Piaractus brachypomus*, and *Utiaritichthys*. It is absent in *Acnodon, Mylesinus, Myleus setiger, Mylossoma, Piaractus mesopotamicus* and *Tometes*. An additional promissing character that can be used in combination with other characters herein proposed to refine the identity of fossil occurrences is the number of cusps on the labial cutting edge; the specimen illustrated by Monsch (1998) presents three cusps, whereas the specimen illustrated byRubilar (1994) presents one or two cusps. This character needs to be further documented across Serrasalmids in order assess its value as a tool for identification of isolated teeth.

> Genus *Piaractus*
>
> *Piaractus* aff. *brachypomus* (Figures 3a-b,d-e)

### Material examined

MUN 37664, two isolated teeth.

### Description

Commissural teeth D4-7, exact serial origin uncertain. Crown well-preserved with remains of cutting edge with tip eroded. Underlying bone preserved in both specimens, showing non-lingually deflected central position of cutting edge. Occlusal flat surface absent.

### Remarks

Commissural teeth D4-7 are distinct from other teeth in the same series in being generally small and with deflected crowns. They show a lingually directed cutting edge that makes them unsuitable for crushing food items, as opposed to teeth D0-3. The fossil specimens herein studied are strongly molariform and with a very low cutting edge that is not lingually deflected, a condition seen in both *Colossoma* and *Piaractus* among serrasalmids. *Colossoma* still has an evident unicuspid cutting edge, a trait far less prominent in *Piaractus*. The presence of unicuspid low cutting edge oberved in these the fossil specimens support their alignment with the genus *Piaractus*. Among the species of this genus the fossil specimens most closely resemble *P. brachypomus*.

> Order Siluriformes
>
> Suborder Loricarioidei
>
> Family Callichthyidae
>
> Gen. sp. *incertae sedis* (Figures 5b-g)

**FIG. 5:**
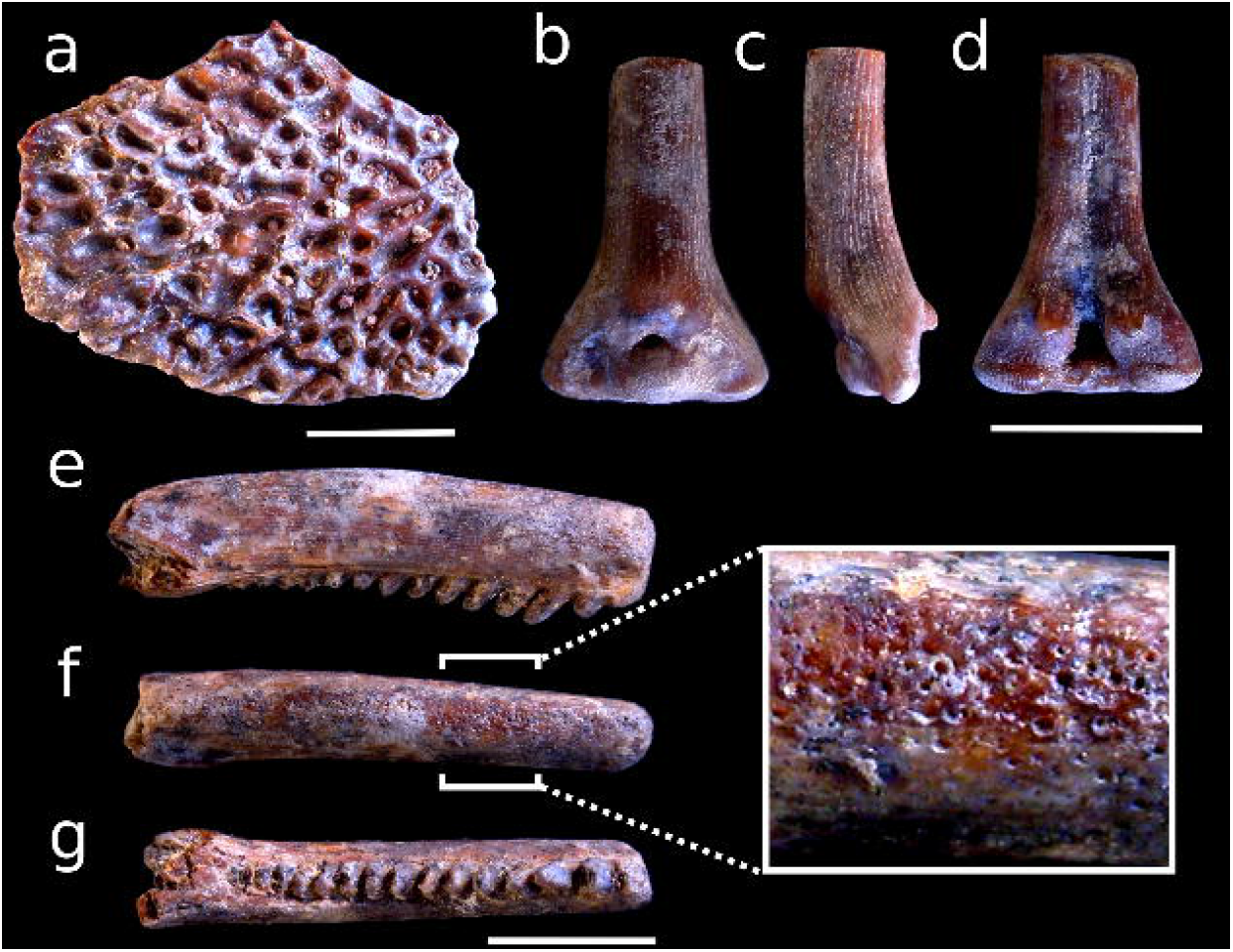
Siluriforms from the middle Castilletes Formation in the locality Makaraipao. a) *Phractocephalus* sp., nuchal plate fragment. b-d) Callichthyidae gen. et sp. indet., dorsal-fin spine fragment in anterior, lateral, and posterior views respectively. e-g) Callichthyidae gen. et sp. indet., pectoral-fin spine fragment in dorsal, anterior, and posterior views respectively. Scale bars equal 10 mm in a, and 5 mm in the remaining sections.

### Material examined

MUN 37803, fragments of one dorsal and one pectoral spines, the former preserving the spine base.

### Description

Dorsal spine with base and proximal portion of shaft. Base triangular in outline with small round base foramen; anterior articular facet inverse-trapezoidal in outline and finely ornamented with vertical ridges. Inflection point of anterior longitudinal ridge poorly developed and tubercle-shaped; anterior longitudinal ridge absent. Anterior fossae wide, oval in outline. Lateral condyles poorly developed, with oval lateral articular surfaces and vertical ridges. Posterior processes shorter than lateral condyles, deflected ventrally. Spine shaft lacking ornaments other than odontode alveoli covering entire anterior surface.

Right pectoral spine represented by shaft from deflection point of dorsal process to about 1/3 of shaft length. Shaft slightly depressed and oval in transverse section. Dorsal and ventral ornaments consisting of parallel, fine ridges; posterior ornament consisting of retrorse compressed blades with dorsal and ventral cutting edges, spanning half of vertical space on posterior surface, increasing in size distally. Fine odontode alveoli covering entire anterior surface of shaft.

### Remarks

Dorsal- and pectoral-fin spines have seldom been identified beyond family in the siluriform fossil record. Callichthyid fin-spine remains have been previously reported in the fossil record. Lundberg (1997) records cf. *Hoplosternum* in the middle Miocene La Venta fauna in central Colombia based on part of the cranium and isolated pectoral spine fragments with odontode alveoli and strong posterior ornaments, straight to slightly retrorse. The combination of odontodes and posterior ornamentation is unique to the Callichthyidae among Loricarioidei.

Spines show strong sexual dimorphism in at least some callichthyid genera. Superficial bone overgrowth of the pectoral spine shaft in males of species of *Callichthys* and *Hoplosternum* obscures ornaments on the posterior surface of the pectoral spine. This does not happen in females and juveniles, which have visible posterior ornaments in the form of straight to slightly retrorse spinules in *Callichthys* and flat spinules in *Hoplosternum*. Thus, the presence or absence of posterior ornamentation in itself cannot be used for identifying taxa. The shape of the spine, on the other hand, is diagnostic for some taxa provided comparisons are restricted to subadults and females. Spine fragments with odontode alveoli but lacking posterior ornament are present in the Loricariidae, thus allowing potential ambiguity with male callichthyids. Loricariids often show a hypertrophied row of odontode alveoli on the dorso-posterior angle of the spine (Ballen & Vari 2012) which are absent in Callichthyids (pers. obs.); this feature aids in distinguishing pectoral spine fragments from those families.

The dorsal spine shaft is very wide in anterior view and antero-posteriorly compressed in species of the genera *Hoplosternum* and *Lepthoplosternum*. This is not seen in the specimen herein studied, which have a regular cross-section outline. The anterior articular facet of the dorsal spine is inversely trapezoid in *Dianema* and *Megalechis*, in contrast to the oval outline in the fossil specimen. The conflicting conditions seen in the fossil specimen preclude any further refinement to the genus level as discussed above.

The Callichthyidae has a fossil record that extends from the late Paleocene to the Pleistocene. Reis (1998) reviewed the fossil record, reassessing the phylogenetic position of *Corydoras revelatus* from the Maiz Gordo Formation in Argentina, the most complete fossil taxon of the family. He also discussed some additional callichthyid fossil occurrences such as *Hoplosternum* sp. from the middle Miocene Honda group in Colombia, and scattered indeterminate remains from the Acre fauna in Brazil and the Pleistocene Luján Formation in Argentina. Lundberg *et al*. (2010) further added an occurrence of cf. *Hoplosternum* from the Madre de Dios fauna without further comments or description of the specimens, they also reported specimens of Callichthyinae and Corydoradinae from the Solimões Formation, Callichthyidae from Madre de Dios, and the previously known *Hoplosternum* sp. from the Honda group. Cione & Baez (2007) reviewed the Argentinian fish fossil record and reported the genera *Corydoras* and *Callichthys* from the Pleistocene of the Bahía Blanca area, also confirming an age of late Paleocene for the Maiz Gordo records of *Corydoras revelatus*. The family Callichthyidae is so far absent in the Venezuelan Urumaco Formation. Thus, the occurrences herein reported are the northernmost fossil record of the family.

> Suborder Siluroidei
>
> Family Pimelodidae
>
> Genus *Phractocephalus*
>
> *Phractocephalus* sp. (Figure 5a)

### Material examined

MUN 37660, fragment of nuchal plate.

### Description

Preserved portion of nuchal plate flat with strong anastomosing ridges on dorsal surface; some isolated tubercles around pits and less frequently on ridges. Preserved bone thickness nearly uniform; margins not preserved and ventral surface moderately eroded.

### Remarks

The presence of strongly reticulate ridged ornament on the nuchal plate is diagnostic for *Phractocephalus* level among South American siluriforms (Lundberg 1997; Lundberg & Aguilera 2003; Aguilera *et al*. 2008; Azpelicueta & Cione 2016; Rincón *et al*. 2016). Although some species of families Ariidae, Doradidae, and Andinichthyidae can have strongly ornamented nuchal plates, their ornament consists of dense tubercles, never reticulating ridges (Aguilera & De Aguilera 2004; Birindelli 2014; Bogan *et al*. 2018). Cranial bones of *Phractocephalus* other than the nuchal plates, also show similar reticulate ornamentation; however all of those, except the opercle, are not mostly flat as in the preserved specimen. The opercular ornament, however, is not as reticulated as in other head bones and its ridges are arranged in radiating pattern from the proximal region of the bone. Other bones (e.g., sphenotic) are partly flat but their ventral surface is as smooth and uniform as the nuchal plate. In sum, evidence preserved in the present specimen clearly indicates that it is a fragment of the nuchal plate of a representative of *Phractocephalus*.

The catfish genus *Phractocephalus* has long been recorded from Neogene sedimentary units in South America. The taxon is represented by a single extant species, *P. hemioliopterus*, and three extinct ones, *P. acreornatus, P. ivy*, and *P. nassi* from Brazil, Argentina, and Venezuela, respectively (Lundberg & Aguilera 2003; Aguilera *et al*. 2008; Azpelicueta & Cione 2016). *Phractocephalus hemioliopterus* is a large, strictly freshwater migratory fish restricted to two cis-Andean drainages, the Amazon and Orinoco (Lundberg & Aguilera 2003; Naranjo & Espinel 2009). At least one fossil species (*P. nassi*) and material of the genus unidentifiable to species (Lundberg 1997) are known from the Neogene of the Magdalena drainage in Colombia. The genus has long been recognized as evidence of past connections between cis- and trans-Andean drainages before separation due to Andean orogeny (Lundberg 1997; Lundberg *et al*. 1998, 2010). The occurrence of *Phractocephalus* in the middle Miocene of the Castilletes Formation confirms its occurrence in the trans-Andean region and supports the hypothesis of drainage connection between trans- and cis-Andean drainages during the middle Miocene.

## FAUNAL SIMILARITY

The seven freshwater fossil fish faunas varied in genus richness. The Urumaco fauna has the richest assemblage with 13 genera; Ituzaingó, Rio Acre, and La Venta show intermediate values with six, eight, and 11 genera respectively, whereas Makaraipao, Contamana, and Fitzcarrald have four genera each. The binary similarity index indicates that the middle Miocene Makaraipao fishes cluster apart from other middle Miocene western Amazonian faunas including La Venta, Fitzcarrald, and Contamana (Figure 6a). The late Miocene pattern is similar, with Urumaco clustering apart from western Amazonia and Parana sites (Figure 6b). Nodal AU support values range between 71 and 94, whereas bootstrap values were consistently lower, ranging from 21 to 61. Observed nodal values of AU are considered as medium to high supporting the clustering structure. Similarity structure does not correlate with geographic linear proximity or mean age of the faunal assemblages (Supplementary section S3).

**FIG. 6:**
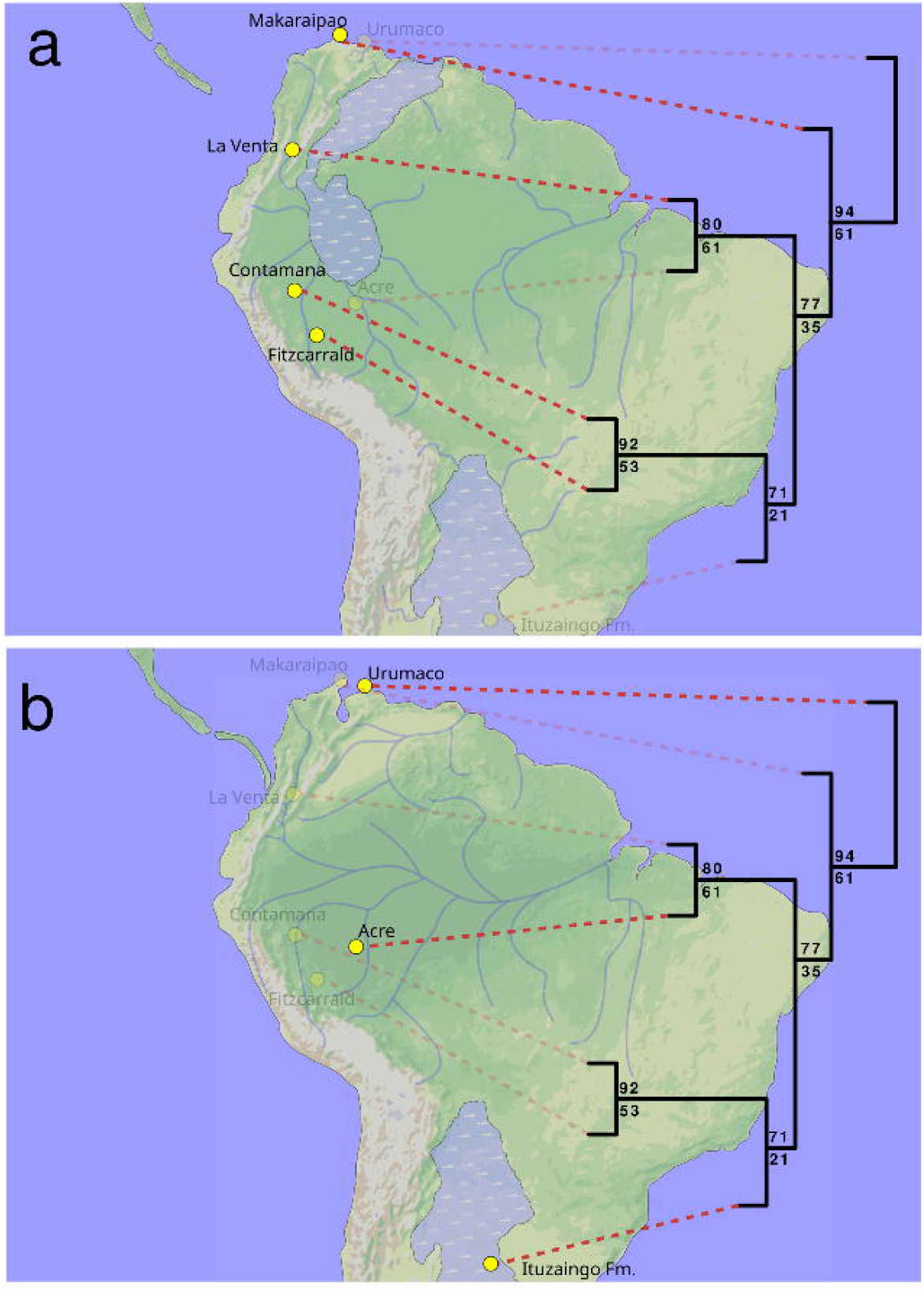
Faunal similarity and in geographic and temporal context. Paleogeographic reconstructions of South America: a) ca. 13 Ma, masking localities that are younger than 11 Ma; b) ca. 8 Ma, shading localities that are older than 11 Ma. The similarity structure does not correlate with mean age of the faunas or with geographic proximity (also see Supplementary section S3). Paleogeographic reconstructions based on Jaramillo *et al*. (2017), Hoorn *et al*. (2010), and Jaramillo et al. (unpubl. data). Values above and below nodes in the dendrogram represent the AU and BT values respectively (Suzuki *et al*. 2019).

The modern faunal similarity cluster recovers two distinct groups, one corresponding to trans-Andean drainages (i.e., Caribbean drainages, Maracaibo, Atrato, Magdalena-Cauca-Sinú) and other cis-Andean comprising mostly Amazonian-Orinocoan-Guyanese basins as well as the southern Paraná Paraguay basin (Figure 7). The cis-Andean cluster comprises an array of basins where the Orinoco drainage (i.e., Apure, upper and lower Orinoco) are a single group clustered within Amazonia, whereas Guiana Shield drainages are intermixed with other Amazonian drainages.

**FIG. 7:**
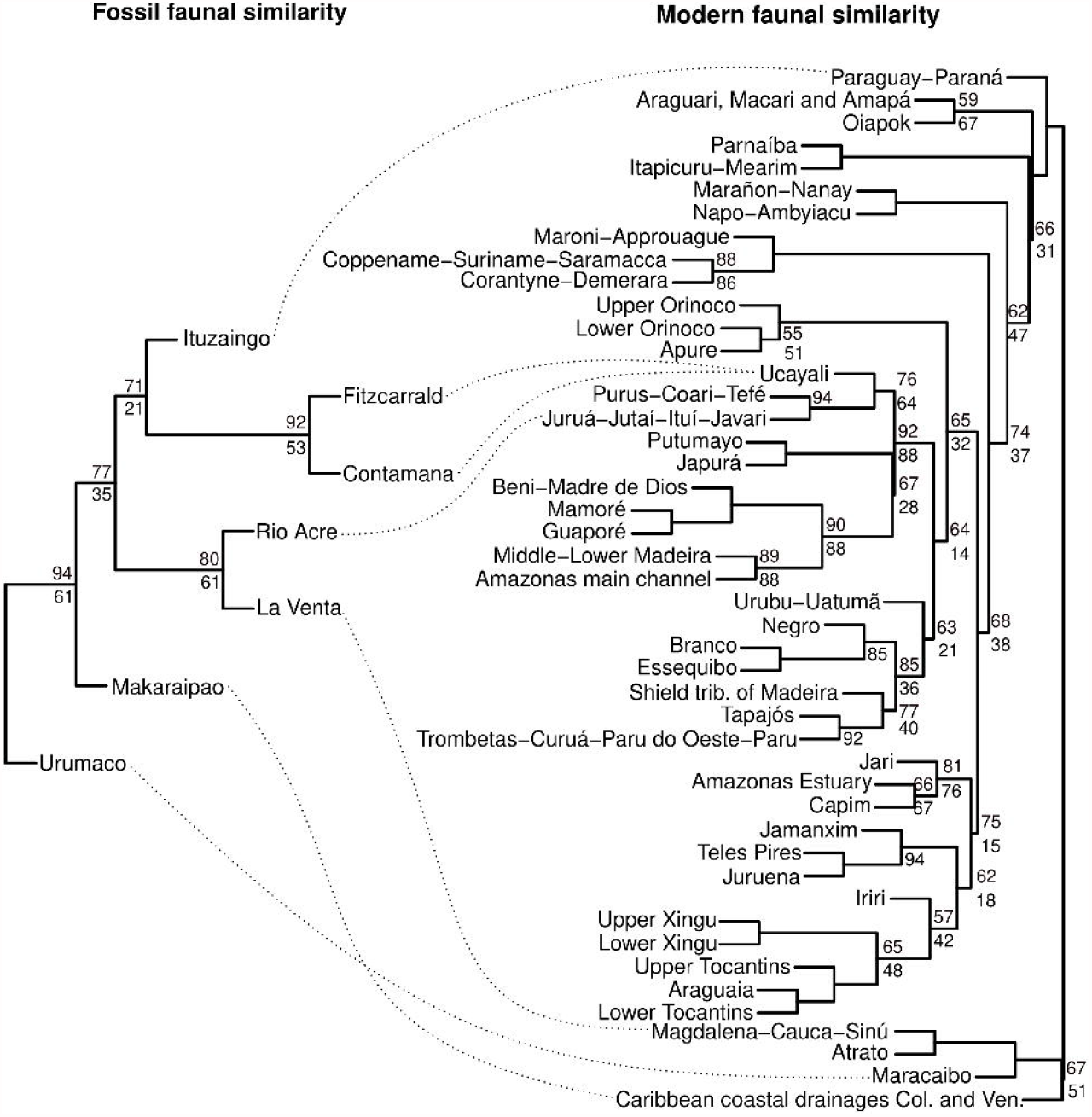
Ancient and modern faunal similarity in South America. Dotted lines match the fossil localities with their corresponding modern drainage. Support values above and below each node are AU and BP respectively, only values lower than 95 are shown. Col. and Ven. equal Colombia and Venezuela respectively, whose coastal drainages are small and numerous which drain the Andes into the Caribbean.

There are two main differences between the extant and Miocene clusters. The extant cis-trans-Andean split is not recovered during the Miocene because the La Venta and Rio Acre basins are clustered together (Figures 6,7). The extant southern Paraná-Paraguay fauna clusters apart from all other Amazonian/Orinoco drainages whereas during the Miocene, the Ituzaingo fauna (that correspond to Paraná) is clustered within Amazonian drainages.

## DISCUSSION

### Paleoecology

The Makaraipao ichthyofauna offers some clues about the middle Miocene paleoenvironmental conditions in the Guajira Peninsula. All of the specimens whose taxonomic identity is at least below the family rank represent extant Amazonian freshwater fish fauna (Figure 4), and none are found in trans-Andean drainages. Large catfishes such as *Phractocephalus* indicate large river channels with long courses permitting reproductive migrations. The only extant species of the genus (*P. hemioliopterus*) migrates distances of ∼300 km (Hahn *et al*. 2019) during the reproductive season; this type of drainage is absent nowadays in the Guajira Peninsula and the coastal drainages in northern Venezuela and the Maracaibo drainages.

*Lepidosiren* is restricted to peripheral, lentic or stagnant environments associated with lagoons and floodplains, indicating that such habitats were also present in Makaraipao (Ballen & Moreno-Bernal 2019). Herbivorous serrasalmids such as *Mylossoma* and *Piaractus* are known for their strong trophic reliance on fruits and seeds from the riparian forest. Their presence suggests that riverine environments in the Guajira Peninsula had gallery forest that provided such resources to the river channel. Lastly, the Callichthyidae is an ecologically very diverse catfish family, so that associated paleoecological parameters cannot at this time be established on the basis of familial assignment only.

Overall, the paleoenvironmental conditions indicated by the fish assemblage indicate year-round rivers with forested vegetation and a surplus of water, which contrasts with the arid and desertic landscape of Guajira nowadays. Similar results were also found by examining the isotopic sclerochronology, which indicates enhanced precipitation (Scholz *et al*. 2020), and the palynological record, which indicates a forested vegetation and high mean annual precipitation (*∼* 2 m/year) (Jaramillo *et al*. 2020).

### Faunal similarity

Continental fossil faunas of Miocene age have long been recognized in South America. Earlier studies based on the fossil mammal assemblage have consistently shown more similarity between the Acre and Urumaco faunas, whereas the older La Venta fauna has been shown to have a distinctive mammal fauna (Cozzuol 2006; Latrubesse *et al*. 2010; Carrillo *et al*. 2015). Cozzuol (2006) found the Acre and Urumaco faunas to be most similar to each other than to the Mesopotamian and the La Venta fauna, while the latter was found to be the most dissimilar of all. Carrillo *et al*. (2015) suggest complex faunal patterns in time and space on the basis of more numerous faunal units and a richer dataset than the earlier study by Cozzuol (2006). Their analyses indicate that the La Venta fauna clusters with the Fitzcarrald fauna and to a lesser degree with Collon Curá, while Urumaco is most similar to Acre and the Mesopotamian in agreement with Cozzuol (2006). According to Carrillo *et al*. (2015) the Mesopotamian includes the fossil assemblage that we herein refer to as the Ituzaingó fish fauna in order to reinforce the stratigraphic and geographic provenance of the fossil association rather than the biochronological implication of the term Mesopotamian, that has a complex application (Cione *et al*. 2000; Cozzuol 2006). Several works refer to the Mesopotamian between quotation marks to acknowledge the complex status of this artificial biochronological name when referring to the faunal association instead (Cione *et al*. 2000; Cozzuol 2006; Latrubesse *et al*. 2010; Brandoni 2011).

However, freshwater fish assemblages suggest a different biogeographic scenario. La Venta clusters with Acre, whereas Urumaco and Makaraipo are dissimilar with respect to all other sites in South America (Figure 6a,b). Urumaco has a large number of taxa not found anywhere else (Lundberg *et al*. 2010).. The components identified in both Contamana and Fitzcarrald are but a small proportion of the recovered fossil material (e.g., supplementary material in Antoine *et al*. 2016), and several specimens await further study; therefore, the similarity between Contamana and Fitzcarrald may change. We found that the pairwise similarity did not correlate with either linear distance (admittedly a poor proxy for geographic relatedness) or mean age of the fossil fauna (Supplementary section S3).

The similarity structure suggests four hydrographic regions: A northern region (Makaraipao in the middle Miocene and Urumaco in the late Miocene), a northwestern region in central Colombia and northwestern Amazonia (La Venta in the middle Miocene and Acre in the late Miocene), a central Amazon region (Contamana and Fitzcarrald, both in the middle Miocene), and a southern region (Ituzaingó, in the late Miocene). However, this pattern is subtle and needs to be confirmed as we improve our knowledge of Amazonian fossil fish assemblages including Acre, Contamana, and Fitzcarrald.

These four regions broadly correspond to the modern similarity structure although there are some differences. The Miocene similarity of La Venta with Rio Acre faunas indicates that this hydrological connection was severed when completion of the eastern Andes uplift in the late Neogene separated Amazonian and trans-Andean drainages. The differences in similarity patterns between the Miocene assemblages and the present-day basins might be due in part to differences in local ecological conditions (e.g., water chemistry; Arbeláez *et al*. 2008; Bogotá-Gregory *et al*. 2020) in addition to zoogeographic patterns in space, or faunal turnover in time. This is a reasonable scenario given that fishes respond to barriers differently than terrestrial organisms (Dagosta & Pinna 2019). Further testing of these alternatives depends on more detailed study of the fossil freshwater fishes of the Contamana and Fitzcarrald faunas.

## CONCLUSION

Fossil vertebrates are key component to to reconstruct past environments, climate, biodiversity, and paleogeography. The middle Miocene Makaraipao freshwater fish indicates a fluvial forest system providing food sources for frugivorous serrasalmids, the presence of flooding systems generating adequate habitats for lepidosirenids, and river channels long and wide enough to support populations of large pimelodid catfishes. Our continental-scale similarity analysis suggest four large paleohydrographic systems during the Neogene with the northern region of South America (Urumaco and Makaraipao) being one of them. Our study highlights several knowledge gaps that will allow us to better understand the past diversity of freshwater fishes in the Neotropics.

## Supporting information

Supplementary section

## ACKNOWLEDGEMENTS

We thank Jaime Escobar and Natalia Hoyos (MUN, Universidad del Norte) for allowing access to fossil specimens from the Castilletes Formation and also for their hospitality during a visit to their institution. Carlos DoNascimiento (IAvH) and Saúl Prada (MPUJ) granted access to collections under their care. John Lundberg and Mark Sabaj (ANSP) allowed us to study the ichthyological collections under their care. The Fundação de Amparo à Pesquisa do Estado de São Paulo (FAPESP) is acknowledged for funding through doctoral scholarships awarded to GAB and FCPD (processes 2014/11558-5 and 2011/23419-1), a BEPE internship awarded to GAB (process 2016/02253-1), and the project “Diversity and Evolution of Gymnotiformes” (process 2016/19075–9). The CNPq is also acknowledged for funding awarded to FCPD (process 405643/2018-7). Part of this research was funded through a Böhlke endowment, and a STRI fellowship, NSF grant EAR-0957679, the National Geographic Society, the Smithsonian Tropical Research Institute, the Anders Foundation, 1923 Fund and Gregory D. and Jennifer Walston Johnson. The Museu de Zoologia da Universidade de São Paulo, Instituto de Ciencias Naturales (Universidad Nacional de Colombia) and Smithsonian Tropical Research Institute (Smithsonian Institution) provided logistical support and workspace during different phases of this work. We thank the 2013 and 2014 field crews of the Guajira Paleontology Project funded by STRI as well as the local Wayuu communities that hosted our field efforts. Carlos DoNascimiento allowed access to photographs of the pectoral spine of *C. oibaensis*. Sandra Reinales provided insightful comments and suggestions during all phases of this study; her help, criticism, and support were key for the completion of this project. GAB thanks Jorge W. Moreno-Bernal, Orangel Aguilera, Annie Hsiou, Maria Camila Vallejo-Pareja, and John Lundberg for discussions on the paleontology of the Neotropics during the Cenozoic. Victor A. Tagliacollo, Gustavo Burin, and Flávio Lima are acknowledged for their insights during the assessment of the doctoral thesis by GAB. Alberto Carvalho and Heraldo Britski provided information on the whereabouts of the type material of *Lepidosiren megalos* and other fossil specimens deposited in MZUSP in the time of Silva Santos. The late Javier Maldonado provided support and access to facilities during an internship awarded to GAB at the Pontificia Universidad Javeriana in 2016. His premature passing did not allow him to see this project completed and we dedicate this paper to his memory.

## AUTHOR CONTRIBUTIONS

**Conceptualization** G Ballen, C Jaramillo; **Data curation** G Ballen, F Dagosta; **Formal analysis** G Ballen; **Funding acquisition** C Jaramillo, G Ballen, M de Pinna; **Investigation** G Ballen, F Dagosta; **Methodology** G Ballen, C Jaramillo; **Project** administration G Ballen; **Resources** C Jaramillo, M de Pinna; **Software** G Ballen; **Supervision** C Jaramillo, M de Pinna; **Validation** G Ballen; **Visualization** G Ballen; **Writing-Original draft preparation** G Ballen; **Writing-Review & editing** G Ballen, C Jaramillo, F Dagosta, and M de Pinna.

## APPENDIX

### COMPARATIVE SPECIMENS

#### Lepidosirenidae

***Lepidosiren paradoxa*: MZUSP 35634, 41101, 50036, 35633**.

#### Serrasalmidae

***Acnodon*** sp.: MZUSP 20361. ***Metynnis fasciatus***: MZUSP 20555. ***Mylesinus schomburgkii***: MZUSP 101455. ***Myleus setiger***: MZUSP 15673. ***Myleus*** sp.: MZUSP 91685. ***Myloplus rhomboidalis***: MZUSP 102136. ***Myloplus schomburgkii***: MZUSP 97639. ***Myloplus torquatus***: MZUSP 95388. ***Mylossoma acanthogaster***: IAvH-P 15900. ***Mylossoma duriventre***: MZUSP 89523. ***Piaractus brachypomus***: MZUSP 20376. ***Piaractus mesopotamicus***: MZUSP 89508. ***Pygopristis denticulata***: MZUSP 57580. ***Tometes kramponhah***: MZUSP 110948. ***Utiaritichthys sennaebragai***: MZUSP 93692

#### Callichthyidae

***Aspidoras albater***: MZUSP 28599, 28, 14.59–39.76; 50157, 1, cs. ***Aspidoras microgaleus***: MZUSP 86842, 1, cs. ***Brochis splendens***: MZUSP 89377, 5, 47.16– 57.47. ***Callichthys callichthys***: MZUSP 43599, 1, 121.12; MZUSP 93043, 8, 47.68– 72.60; MZUSP 109087, 2, 116.71–145.12; MZUSP 84199, 2, cs. ***Callichthys serralabiatum***: MZUSP 93168, 1, 104.65. ***Corydoras araguaiensis***: MZUSP 86269, 1, cs. ***Corydoras ehrhardti***: MZUSP 81572. ***Corydoras*** cf. ***guianensis***: MZUSP 107151, 2, cs. ***Dianema*** sp.: MZUSP 30862, 2, cs. ***Hoplosternum littorale***: MZUSP 85987, 1, 84.91; MZUSP 94658, 1, 95.46; MZUSP 117107, 1, cs. ***Lepthoplosternum pectorale***: MZUSP 83608, 2, cs; MZUSP 112403, 1, 57.13. ***Megalechis thorocata***: MZUSP 25451, 2, cs. ***Scleromystax barbatus***: MZUSP 37723, 2, cs.

#### Pimelodidae

***Bergiaria westermanni***: MZUSP 85627. ***Brachyplatystoma capapretum***: MZUSP 53262, ANSP 178524, 179733, 178524. ***Brachyplatystoma juruense***: ANSP 178514-1,3,4,6, DUF 1071. ***Brachyplatystoma platynemum***: DUF 993, 1076, ANSP uncat, 187321. ***Brachyplatystoma rousseauxii***: ANSP 179794, 179793, 179233, DUF 1051, 981, 1078. ***Brachyplatystoma vaillanti***: ANSP 179799, 178525, 179474, DUF 994, 1164. ***Brachyplaystoma filamentosum***: DUF 1079, 1080, ANSP 179776. ***Calophysus macropterus***: DUF 1199, 1049, 403, ANSP 199813, 178164, 178260, ICNMHN uncat. ***Duopalatinus emarginatus***: MZUSP 85622. ***Exallodontus aguanai***: ANSP 18947, uncat. ***Hemisorubim platyrhynchos***: ANSP uncat, 179234, ICNMHN 7909, MZUSP 7009. ***Hypophthalmus*** sp. “curved”: ANSP uncat. ***Hypophthalmus*** sp. “straight”: ANSP 180993, 178512, 180993, 187103. ***Iheringichthys labrosus***: ANSP 180505, MZUSP 78459, 25102. ***Leiarius marmoratus***: MZUSP 108333. ***Leiarius perruno***: ANSP uncat-I, uncat-II, ICNMHN 2158. ***Leiarius pictus***: MZUSP 82577. ***Leiarius*** sp.: ANSP 178526, 178527, uncat without skull, DUF 1054, 1056, 1036, 1055, 1037. ***Luciopimelodus pati***: ANSP 178798, MZUSP 78464, 78457. *Megalonema platanum*: MZUSP 78465. ***Megalonema platycephalum***: ANSP 179249, 178515. ***Megalonema*** sp.: MZUSP 92604. ***Parapimelodus nigribarbus***: MZUSP 78451. ***Parapimelodus valenciennesi***: MZUSP 78466, ANSP 178800.***Phractocephalus hemioliopterus***: ANSP 179559, 179553, 179554, ICN uncat. ***Pimelodina flavipinnis***: ANSP uncat, 178513, 178516. ***Pimelodus argenteus***: ANSP 181017, uncat. ***Pimelodus fur***: MZUSP 22566. ***Pimelodus grosskopfii***: ICNMHN 6867. ***Pimelodus maculatus***: MZUSP 85486, 110379. ***Pimelodus microstoma***: MZUSP 22696, 22712 (paratype of ***Pimelodus heraldoi***). ***Pimelodus mysteriosus***: ANSP 180506, MZUSP 90595. ***Pimelodus ortmanni***: MZUSP 50053. ***“Pimelodus” altipinnis***: MZUSP 58328. ***“Pimelodus” ornatus***: ANSP 178452, 180985, MZUSP 34480, 109128. ***Pinirampus argentinus***: ANSP 181016. ***Pinirampus pirinampu***: ANSP uncat, 178530. ***Platynematichthys notatus***: ANSP uncat, 178528. ***Platysilurus malarmo***: ANSP uncat., iavh-p 11815, 11819, 11894 ***Platysilurus mucosus***: DUF 986, uncat, ANSP 178508, 178509, iavh-p 17638, 10850, 10956, 6006 ***Platystomatichthys sturio***: ANSP uncat, SU 22463, ICNMHN 15201. ***Platystomatichthys*** sp. “Xingu”: MZUSP 58364. ***Propimelodus*** sp.: ANSP 180939. ***Pseudoplatystoma corruscans***: ANS 188913, MZUSP 78477. ***Pseudoplatystoma fasciatum***: ANSP 177346. ***Pseudoplatystoma magdaleniatum***: ICNMHN 6860. ***Pseudoplatystoma metense***: ANSP 149541. ***Pseudoplatystoma reticulatum***: ANSP 188912. ***Pseudoplatystoma*** sp.: DUF 1125, ICNMHN uncat. ***Pseudoplatystoma tigrinum***: DUF 921, uncat, ANSP 187010. ***Sorubim cuspicaudus***: MBUCV uncat, DUF 932. ***Sorubim elongatus***: ICNMHN 15039. ***Sorubim lima***: ANSP 178507. ***Sorubim trigonocephalus***: ANSP 188824. ***Sorubimichthys planiceps***: ANSP 179235, 17850. ***Steindachneridion scripta***: MZUSP 78463. ***Zungaro zungaro***: ANSP uncat, DUF 982, ICN uncat, MPUJ 13213, MZUSP 96290, 108335, 94859, 96188, 151727.

## BIBLIOGRAPHY

Agnolin, F. L. 2010. A new species of the genus Atlantoceratodus (Dipnoiformes: Ceratodontoidei) from the Uppermost Cretaceous of Patagonia and a brief overview of fossil dipnoans from the Cretaceous and Paleogene of South America. Brazilian Geographical Journal: Geosciences and Humanities Research Medium, 1, 162–210.

Aguilera, O., Oliveira, G. A. S., Lopes, R. T., Machado, A. S., Santost. M. Dos, Marques, G., Bertucci, T., Aguiar, T., Carrillo-Briceño, J. D., Rodriguez, F. and Jaramillo, C. 2017. Neogene Proto-Caribbean porcupinefishes (Diodontidae). PLoS ONE, 12, 1–26.

Aguilera, O. A. and De Aguilera, D. R. 2004. Amphi-American neogene sea catfishes (Siluriformes, Ariidae) from northern south America. Special Papers in Palaeontology, 71, 29–48.

Aguilera, O. A., Bocquentin, J., Lundberg, J. G. and Maciente, A. 2008. A new cajaro catfish (Siluriformes: Pimelodidae: Phractocephalus) from the Late Miocene of southwestern Amazonia and its relationship to Phractocephalus nassi of the Urumaco Formation. Palaeontologische Zeitschrift, 82, 231–245.

Aguilera, O. A., Moraes-Santos, H., Costa, S. A. R. F., Ohe, F., Jaramillo, C. A. and Nogueira, A. 2013a. Ariid sea catfishes from the coeval Pirabas (Northeastern Brazil), Cantaure, Castillo (Northwestern Venezuela), and Castilletes (North Colombia) formations (early Miocene), with description of three new species. Swiss Journal of Palaeontology, 132, 45–68.

Aguilera, O. A., Lundberg, J. G., Birindelli, J. L. O., Sabaj Pérez, M. H., Jaramillo, C. A. and Sánchez-Villagra, M. R. 2013b. Palaeontological evidence for the last temporal occurrence of the ancient Western Amazonian river outflow into the Caribbean. PLoS ONE, 8, 1–17.

Albert, J. S. and Reis, R. E. 2011. Historical Biogeography of Neotropical Freshwater Fishes. In Albert, J. S.and Reis, R. E. (eds.) University of California Press, 388 pp.

Almeida-Val, V. M. F., Nozawa, S. R., Lopes, N. P., Aride, P. H. R., Mesquita-Saad, L. S., Silva M. de N. P. Da, Honda, R. T., Ferreira-Nozawa, M. S. and Val, A. L. 2011. Biology of the South American Lungfish, Lepidosiren paradoxa. In Jorgensen, J. M.and Joss, J. (eds.) The Biology of Lungfishes, CRC Press, Boca Raton, 129–147 pp.

Alves, Y. M., Machado, L. P., Paglarelli Bergqvist, L. and Brito, P. M. 2013. Redescription of two lungfish (Sarcopterygii: Dipnoi) tooth plates from the Late Cretaceous Bauru Group, Brazil. Cretaceous Research, 40, 243–250.

Amson, E., Carrillo, J. D. and Jaramillo, C. A. 2016. Neogene sloth assemblages (Mammalia, Pilosa) of the Cocinetas Basin (La Guajira, Colombia): implications for the Great American Biotic Interchange. Palaeontology, 59, 563–582.

Andrade, M. C., Ota, R. P., Bastos, D. A. and Jégu, M. 2016. A new large Myloplus Gill 1896 from rio Negro basin, Brazilian Amazon (Characiformes: Serrasalmidae). Zootaxa, 4205, 571–580.

Antoine, P.-O., Abello, M. A., Adnet, S., Altamirano Sierra, A. J., Baby, P., Billet, G., Boivin, M., Calderón, Y., Candela, A., Chabain, J., Corfu, F., Croft, D. A., GanerØd, M., Jaramillo, C. A., Klaus, S., Marivaux, L., Navarrete, R. E., Orliac, M. J., Parra, F., Pérez, M. E., Pujos, F., Rage, J. C., Ravel, A., Robinet, C., Roddaz, M., Tejada-Lara, J. V., Vélez-Juarbe, J., Wesselingh, F. P. and Salas-Gismondi, R. 2016. A 60-million-year Cenozoic history of western Amazonian ecosystems in Contamana, eastern Peru. Gondwana Research, 31, 30–59.

Arbeláez, F., Duivenvoorden, J. F. and Maldonado-Ocampo, J. A. 2008. Geological differentiation explains diversity and composition of fish communities in upland streams in the southern Amazon of Colombia. Journal of Tropical Ecology, 24, 505–515.

Azpelicueta, M. de las M. and Cione, A. L. 2016. A southern species of the tropical catfish genus Phractocephalus (Teleostei: Siluriformes) in the Miocene of South America. Journal of South American Earth Sciences, 67, 221–230.

Ballen, G. A. and Vari, R. P. 2012. Review of the Andean armored catfishes of the genus Dolichancistrus Isbrücker (Siluriformes: Loricariidae). Neotropical Ichthyology, 10, 499–518.

Ballen, G. A. and Moreno-Bernal, J. W. 2019. New records of the enigmatic Neotropical fossil fish Acregoliath rancii (Teleostei incertae sedis) from the middle Miocene Honda group of Colombia. Ameghiniana, 56, 431–440.

Ballen, G. A. and PINNA M.C.C. De. 2021. A standardized terminology of spines in the order Siluriformes (Actinopterygii: Ostariophysi). Zoological Journal of the Linnean Society, 1–25.

Birindelli, J. L. O. 2014. Phylogenetic relationships of the South American Doradoidea (Ostariophysi: Siluriformes). Neotropical Ichthyology, 12, 451–564.

Bogan, S., Agnolin, F. L. and Scanferla, A. 2018. A new Andinichthyidae catfish (Ostariophysi, Siluriformes) from the Paleogene of northwestern Argentina. Journal of Vertebrate Paleontology, 38.

Bogan, S., Sidlauskas, B., Vari, R. P. and Agnolin, F. L. 2012. Arrhinolemur scalabrinii Ameghino, 1898, of the late Miocene - a taxonomic journey from the Mammalia to the Anostomidae (Ostariophysi: Characiformes). Neotropical Ichthyology, 10, 555–560.

Bogotá-Gregory, J. D., Lima, F. C. T., Correa, S. B., Silva-Oliveira, C., Jenkins, D. G., Ribeiro, F. R., Lovejoy, N. R., Reis, R. E. and Crampton, W. G. R. 2020. Biogeochemical water type influences community composition, species richness, and biomass in megadiverse Amazonian fish assemblages. Nature Scientific Reports, 10, 1–15.

Brandoni, D. 2011. The Megalonychidae (Xenarthra, Tardigrada) from the late Miocene of Entre Ríos Province, Argentina, with remarks on their systematics and biogeography. Geobios, 44, 33–44.

Cadena, E. and Jaramillo, C. A. 2015a. The first fossil skull of Chelus (Pleurodira: Chelidae, Matamata turtle) from the early Miocene of Colombia. Palaeontologica Electronica, 18.2, 1–10.

Cadena, E. and Jaramillo, C. A. 2015b. Early to Middle Miocene Turtles from the Northernmost Tip of South America: Giant Testudinids, Chelids, and Podocnemidids from the Castilletes Formation, Colombia. Ameghiniana, 52, 188–203.

Carrillo, J. D., Forasiepi, A., Jaramillo, C. and Sánchez-Villagra, M. R. 2015. Neotropical mammal diversity and the great American biotic interchange: Spatial and temporal variation in South America’s fossil record. Frontiers in Genetics, 5, 1–11.

Carrillo-Briceño, J. D., Luz, Z., Hendy, A., Kocsis, L., Aguilera, O. and Vennemann, T. 2019. Neogene Caribbean elasmobranchs: Diversity, paleoecology and paleoenvironmental significance of the Cocinetas Basin assemblage (Guajira Peninsula, Colombia). Biogeosciences, 16, 33–56.

Cione, A. L., Azpelicueta, M. De Las M., Bond, M., Carlini, A. A., Casciotta, J. R., Cozzuol, M. A., FUENTE M. De La, Gasparini, Z., Goin, F. J., Noriega, J., Scillato-Yané, G. J., Soibelzon, L., Tonni, E. P., Verzi, D. and Vucetich, M. G. 2000. The Miocene vertebrates from Paraná, eastern Argentina Miocene vertebrates from Entre Ríos province, eastern Argentina. Correlación Geológica, 14, 191–237.

Cione, A. L. and Baez, A. M. 2007. Peces, anfibios e invertebrados cenozoicos de Argentina: los últimos cincuenta años. Asociación Paleontológica Argentina. Publicación Especial, 11, 195–220.

Cione, A. L. and Azpelicueta, M. De Las M. 2013. The first fossil species of Salminus, a conspicuous South American freshwater predatory fish (Teleostei, Characiformes), found in the Miocene of Argentina. Journal of Vertebrate Paleontology, 33, 1051–1060.

Cione, A. L., Dahdul, W. M., Lundberg, J. G. and Machado-Allison, A. 2009. Megapiranha paranensis, a new genus and species of Serrasalmidae (Characiformes, Teleostei) from the upper Miocene of Argentina. Journal of Vertebrate Paleontology, 29, 350–358.

Cozzuol, M. A. 2006. The Acre vertebrate fauna: Age, diversity, and geography. Journal of South American Earth Sciences, 21, 185–203.

Criswell, K. E. 2015. The comparative osteology and phylogenetic relationships of African and South American lungfishes (Sarcopterygii: Dipnoi). Zoological Journal of the Linnean Society, 174, 801–858.

Dagosta, F. C. P. and Pinnam. C. C. De. 2019. The fishes of the Amazon: Distribution and biogeographical patterns, with a comprehensive list of species. Bulletin of the American Museum of Natural History, 431, 1–163.

Dahdul, W. M. 2004. Fossil Serrasalmine fishes (Teleostei: Characiformes) from the Lower Miocene of north-eastern Venezuela. Special Papers in Palaeontology, 71, 23–28.

Diaz De Gamero, M.L. 1996. The changing course of the Orinoco River during the Neogene: A review. Palaeogeography, Palaeoclimatology, Palaeoecology, 123, 385–402.

Forasiepi, A. M., Soibelzon, L. H., Suárez Gomez, C., Sánchez, R., Quiroz, L. I., Jaramillo, C. A. and Sánchez-Villagra, M. R. 2014. Carnivorans at the Great American Biotic Interchange: new discoveries from the northern neotropics. Naturwissenschaften, 101, 965–74.

Gayet, M., Marshall, L. G., Sempere, T., Cappetta, H. and Rage, J. 2001. Middle Maastrichtian vertebrates (fishes, amphibians, dinosaurs and other reptiles, mammals) from Pajcha Pata (Bolivia). Biostratigraphic, palaeoecologic and palaeobiogeographic implications. Palaeogeography, Palaeoclimatology, Palaeoecology, 169, 39–68.

Gelfo, J. N., Goin, F. J., Woodburne, M. O. and De Muizon, C. 2009. Biochronological relationships of the earliest South American paleogene mammalian faunas. Palaeontology, 52, 251–269.

Gregory-Wodzicki, K. M. 2000. Uplift history of the Central and Northern Andes: A review. Geological Society of America Bulletin, 112, 1091–1105.

Hahn, L., Martins, E. G., Nunes, L. D., Câmara, L.F., Da Machado, L. S. and Garrone-Neto, D. 2019. Biotelemetry reveals migratory behaviour of large catfish in the Xingu River, Eastern Amazon. Nature Scientific Reports, 9, 1–15.

Hendy, A. J. W., Jones, D. S., Moreno, F., Zapata, V. and Jaramillo, C. A. 2015. Neogene molluscs, shallow marine paleoenvironments, and chronostratigraphy of the Guajira Peninsula, Colombia. Swiss Journal of Palaeontology, 134, 45–75.

Hoorn, C., Wesselingh, F. P., STEEGEH. Ter, Bermudez, M., Mora, A., Sevink, J., Sanmartín, I., Sanchez-Meseguer, A., Anderson, C. L., Figueiredo, J. P., Jaramillo, C., Riff, D., Negri, F. R., Hooghiemstra, H., Lundberg, J. G., Stadler, T., Särkinen, T. and Antonelli, A. 2010. Amazonia through time: Andean uplift, climate change, landscape evolution, and biodiversity. Science, 330, 927–931.

Jaramillo, C., Sepulchre, P., Cardenas, D., Correa-Metrio, A., Moreno, J. E., Trejos, R., Vallejos, D., Hoyos, N., Martínez, C., Carvalho, D., Escobar, J., Oboh-Ikuenobe, F., Prámparo, M. B. and Pinzón, D. 2020. Drastic vegetation change in the Guajira Peninsula (Colombia) during the Neogene. Paleoceanography and Paleoclimatology, 1–36.

Jaramillo, C., Romero, I., D’Apolito, C., Bayona, G., Duarte, E., Louwye, S., Escobar, J., Luque, J., Carrillo-Briceño, J. D., Zapata, V., Mora, A., Schouten, S., Zavada, M., Harrington, G., Ortiz, J. and Wesselingh, F. P. 2017. Miocene flooding events of western Amazonia. Science Advances, 3, 1–12.

Jégu, M. and, SANTOSG. M. Dos. 1990. Description d’Acnodon senai n. sp. du Rio Jari (Brésil, Amapà) et redescription d’A. normani (Teleostei, Serrasalmidae). Cybium, 14, 187–206.

Kolmann, M. A., Huie, J. M., Evans, K. and Summers, A. P. 2018. Specialized specialists and the narrow niche fallacy: A tale of scale-feeding fishes. Royal Society Open Science, 5, 1–14.

Latrubesse, E. M., Cozzuol, M. A., Silva-Caminha, S.A.F., Da Rigsby, C. A., Absy, M. L. and Jaramillo, C. A. 2010. The Late Miocene paleogeography of the Amazon Basin and the evolution of the Amazon River system. Earth-Science Reviews, 99, 99–124.

López-Fernández, H. and Albert, J. S. 2011. Paleogene radiations. In Albert, J. S. and Reis, R. E. (eds.) Historical Biogeography of Neotropical Freshwater Fishes, University of California Press, Berkeley, CA, 105–117 pp.

Lundberg, J. G. 1997. Freshwater fishes and their paleobiotic implications. In Kay, R. F., Madden, R. H., Cifelli, R. L.and Flynn, J. J. (eds.) Vertebrate Paleontology in the Neotropics: The Miocene Fauna of La Venta, Colombia, Smithsonian Institution Press, 67–91 pp.

Lundberg, J. G. and Aguilera, O. A. 2003. The late Miocene Phractocephalus catfish (Siluriformes: Pimelodidae) from Urumaco, Venezuela: additional specimens and reinterpretation as a distinct species. Neotropical Ichthyology, 1, 97–109.

Lundberg, J. G., Sabaj Pérez, M. H., Dahdul, W. M. and Aguilera, O. A. 2010. The Amazonian Neogene fish fauna. In Hoorn, C. and Wesselingh, F. P. (eds.) Amazonia, Landscape and Species Evolution: A Look into the Past, Blackwell Publishing, 281–301 pp.

Lundberg, J. G., Marshall, L. G., Guerrero, J., Horton, B., Malabarba, M. C. S. L. and Wesselingh, F. 1998. The stage for Neotropical Fish diversification: A history of Tropical South American rivers. In Malabarba, L. R., Reis, R. E., Vari, R. P., Lucena, Z. M.and Lucena, C. A. S. (eds.) Phylogeny and Classification of Neotropical Fishes, Edipucrs, Porto Alegre, 13–48 pp.

Mirande, J. M. 2010. Phylogeny of the family Characidae (Teleostei: Characiformes): from characters to taxonomy. Neotropical Ichthyology, 8, 385–568.

Monsch, K. A. 1998. Miocene fish faunas from the northwestern Amazonia basin (Colombia, Peru, Brazil) with evidence of marine incursions. Palaeogeography, Palaeoclimatology, Palaeoecology, 143, 31–50.

Montes, C., Silva, C. A., Bayona, G., Villamil, R., Stiles, E., Rodriguez-Corcho, A. F., Beltran-Triviño, A., Lamus, F., Muñoz-Granados, M. D., Pérez-Angel, L. C., Hoyos, N., Gomez, S., Galeano, J. J., Romero, E., Baquero, M., Cardenas-Rozo, A. L. and QUADT A. Von. 2021. A Middle to late Miocene trans-Andean portal: Geologic record in the Tatacoa Desert. Frontiers in Earth Science, 8, 1–19.

Moreno, F., Hendy, A. J. W., Quiroz, L., Hoyos, N., Jones, D. S., Zapata, V., Zapata, S., Ballen, G. A., Cadena, E., Cárdenas, A. L., Carrillo-Briceño, J. D., Carrillo, J. D., Delgado-Sierra, D., Escobar, J., Martínez, J. I., Martínez, C., Montes, C., Moreno, J., Pérez, N., Sánchez, R., Suárez, C., Vallejo-Pareja, M. C. and Jaramillo, C. A. 2015. Revised stratigraphy of Neogene strata in the Cocinetas Basin, La Guajira, Colombia. Swiss Journal of Palaeontology, 134, 5–43.

Moreno-Bernal, J. W., Head, J. and Jaramillo, C. A. 2016. Fossil crocodilians from the High Guajira Peninsula of Colombia: Neogene faunal change in northernmost South America. Journal of Vertebrate Paleontology, 36, e1110586.

Naranjo, L. G. and Espinel, J. D. A. 2009. Plan nacional de las especies migratorias. Diagnóstico e identificacion de acciones para la conservacion y el manejo sostenible de las especies migratorias de la biodiversidad en Colombia. Ministerio de Medio Ambiente de Colombia, Bogotá, 1–214 pp.

Oksanen, J., Blanchet, F. G., Friendly, M., Kindt, R., Legendre, P., Mcglinn, D., Minchin, P. R., O’hara, R. B., Simpson, G. L., Solymos, P., Stevens, M. H. H., Szoecs, E. and Wagner, H. 2019. vegan: Community Ecology Package.

Pebesma, E. 2018. Simple features for R: Standardized support for spatial vector data. R Journal, 10, 439–446.

QGIS DEVELOPMENT TEAM. 2019. QGIS Geographic Information System. Open Source Geospatial Foundation Project.

R CORE DEVELOPMENT TEAM. 2018. R: A language and environment for statistical computing.

Reis, R. E. 1998. Systematics, biogeography, and tyhe fossil record of the Callichthyidae: A review of the available data.In Malabarba, L. R., Reis, R. E., Vari, R. P., Lucena, Z. M. S. D.and, LUCENAC. A. S. De (eds.) Phylogeny and Classification of Neotropical Fishes, Edipucrs, Porto Alegre, 351–362 pp.

Rincón, A. D., Solórzano, A., Macsotay, O., Mcdonald, H. G. and Núñez-Flores, M. 2016. A new Miocene vertebrate assemblage from the Río Yuca Formation (Venezuela) and the northernmost record of typical Miocene mammals of high latitude (Patagonian) affinities in South America. Geobios, 49, 395– 405.

Rubilar, A. 1994. Diversidad ictiológica en depósitos continentales miocenos de la Formación Cura-Mallín, Chile (37-39S): Implicancias paleogeográficas. Revista Geologica de Chile, 21, 3–29.

Schaefer, S. A. 2011. The Andes. Riding the Tectonic Uplift. In Albert, J. S.and Reis, R. E. (eds.) Historical Biogeography of Neotropical Freshwater Fishes, University of California Press, 259–278 pp.

Scholz, S. R., Petersen, S. V., Escobar, J., Jaramillo, C., Hendy, A. J. W., Allmon, W. D., Curtis, J. H., Anderson, B. M., Hoyos, N., Restrepo, J. C. and Perez, N. 2020. Isotope sclerochronology indicates enhanced seasonal precipitation in northern South America (Colombia) during the Mid-Miocene Climatic Optimum. Geology, 48, 668–672.

Silva Santos, R. 1987. Lepidosiren megalos n. sp. do Terciario do Estado do Acre; Brasil. Anais da Academia Brasileira de Ciencias, 59, 375–384.

Suarez, C., Forasiepi, A. M., Goin, F. J. and Jaramillo, C. A. 2016. Insights into the Neotropics Prior to the Great American Biotic Interchange: New evidence of mammalian predators from the Miocene of Northern Colombia. Journal of Vertebrate Paleontology, 36, e1029581.

Suzuki, R., Terada, Y. and Shimodaira, H. 2019. pvclust: Hierarchical Clustering with P-Values via Multiscale Bootstrap Resampling.

Tejada-Lara, J. V., Salas-Gismondi, R., Pujos, F., Baby, P., Benammi, M., Brusset, S.,, Frenceschid. D. de, Espurt, N., Urbina, M. and Antoine, P.-O. 2015. Life in proto-Amazonia: Middle Miocene Mammals from the Fitzcarrald Arch (Peruvian Amazonia). Palaeontology, 58, 341–378.

