## Supplementary section for "A fossil fish assemblage from the middle Miocene of the Cocinetas Basin, northern Colombia"

Supplementary material

Gustavo A. Ballen, Carlos Jaramillo, Fernando C. P. Dagosta & Mario C. C. de Pinna

**S3 Correlates to community similarity** **9**

**S4 Raw occurrence data** **13**

```
## Loading required package: permute

## Loading required package: lattice

## This is vegan 2.5-7

## Linking to GEOS 3.9.1, GDAL 3.3.0, PROJ 8.0.1
```

#### S1 Faunal similarity: Expanded analysis

```
### faunal similarity analyses for the Marakaipao fish fauna
dataset <- read.delim(file = "neogeneFishOccs.tab", stringsAsFactors = FALSE)

#################### only freswater taxa
dataset <- dataset[which(dataset$Environment == "Freshwater" |
    dataset$Environment == "Both"), ]

### complete analysis without filtering localities
wholeMatrix <- dataset[, -c(1, 3, 19)]
rownames(wholeMatrix) <- wholeMatrix$Taxon
wholeMatrix <- wholeMatrix[-1]
wholeMatrix <- t(wholeMatrix)

### include only faunas with positive number of occurrences
wholeMatrix <- wholeMatrix[apply(X = wholeMatrix, MARGIN = 1,
    FUN = sum) > 0, ]
```

```r
### number of occurrences per fauna
sort(x = apply(X = wholeMatrix, MARGIN = 1, FUN = sum), decreasing = TRUE)
```

```
##        Urumaco        La.Venta       Rio.Acre      Ituzaingo     Makaraipao
##             13              11              8              6              4
## Solimoes.Pebas    Fitzcarrald      Contamana       Castillo  Loyola.Mangan
##              4               4              4              3              2
##       Rio.Yuca        Utuquina
##              2               1
```

```r
### calculate the distance matrix using Bray-Curtis' method
distMatrixBinarywhole <- pvclust::pvclust(t(wholeMatrix), method.hclust = "average",
    method.dist = "binary")
```

```
## Bootstrap (r = 0.48)... Done.
## Bootstrap (r = 0.59)... Done.
## Bootstrap (r = 0.67)... Done.
## Bootstrap (r = 0.78)... Done.
## Bootstrap (r = 0.89)... Done.
## Bootstrap (r = 1.0)... Done.
## Bootstrap (r = 1.07)... Done.
## Bootstrap (r = 1.19)... Done.
## Bootstrap (r = 1.3)... Done.
## Bootstrap (r = 1.37)... Done.
```

```r
plot(distMatrixBinarywhole, main = "Faunal similarity")
```

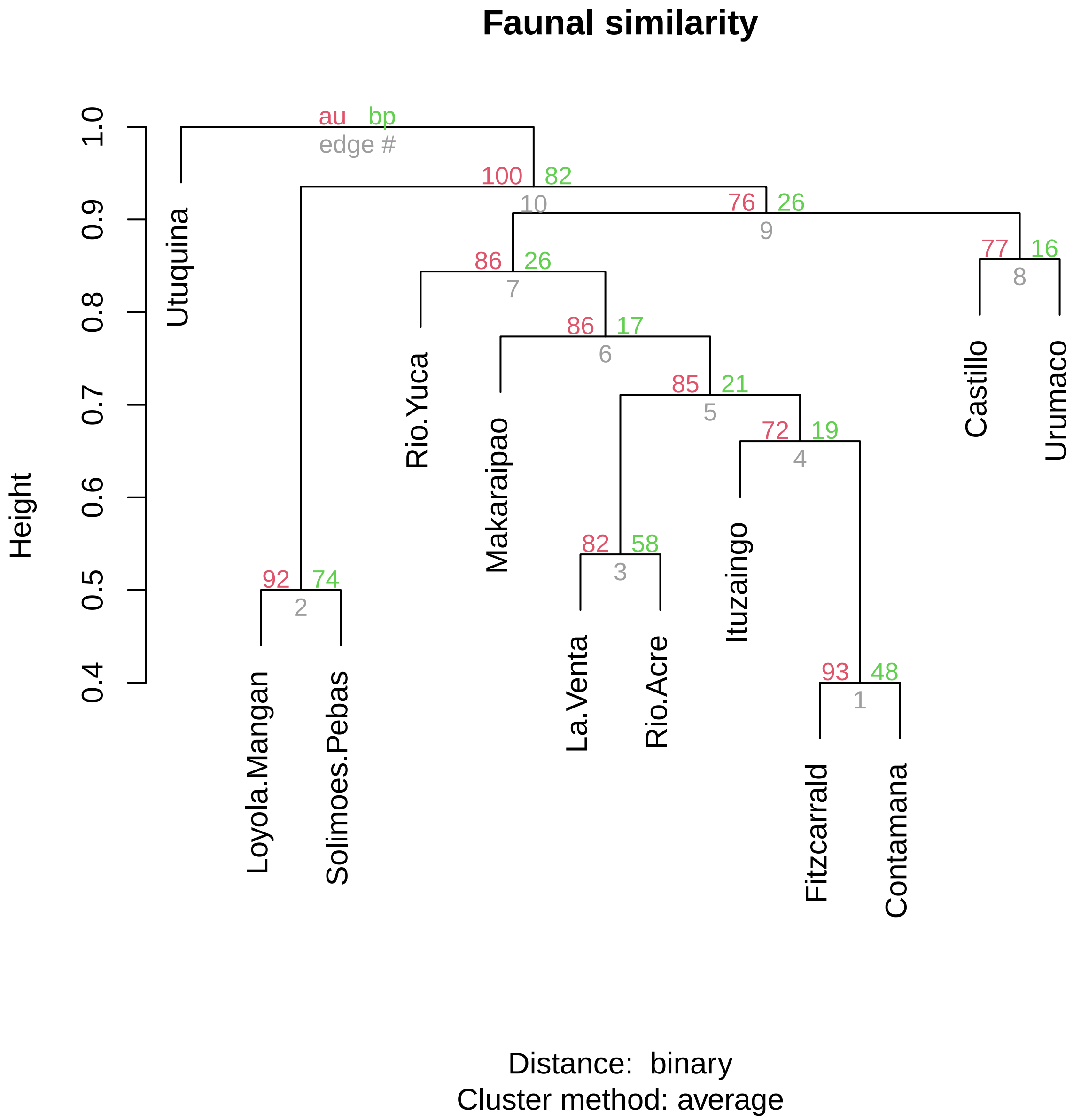

#### S2 Dataset cleaning

Data from Species Link were downloaded and stored for the whole of Chordata, so that specific names could be sliced with the following scripts:

```
### slicer
#!/usr/bin/env bash

# TAXON and FILE are mandatory arguments for the expression (taxon name) TAXON to be
# serched for by grep in the file FILE.
# the result is written to a new file preserving the original FILE filename and
# appending a .out string so it can be differentiated
TAXON=$1
FILE=$2
OUTPUT=`(basename $FILE)`.out
head -n 1 $FILE > $OUTPUT
```

```
grep $TAXON $FILE >> $OUTPUT
NLINES=`wc -l *out | awk '{print $1}'`
echo "Finished writing $OUTPUT with $NLINES lines"
```

```
### speciesLinkPicker.sh
# this script uses slicer, a small program in bash that I wrote for that specific purpose.
# SPLINKFILE is the path to the specieslink file directory
SPLINKFILE=speciesLink_all_49771_20190521013813.txt

# Callichthyidae
slicer Callichthyidae $SPLINKFILE
cp *out callichthyidae/
rm *out

# Lepidosiren
slicer Lepidosiren $SPLINKFILE
cp *out lepidosiren/
rm *out

# Phractocephalus
slicer Phractocephalus $SPLINKFILE
cp *out phractocephalus/
rm *out

# Serrasalmidae
slicer Serrasalmidae $SPLINKFILE
cp *out serrasalmidae/
rm *out
```

#### S2.1 Specific data cleaning

##### S2.1.1 Callichthyidae

```
gbifData <- read.delim("occurrence.txt", header = TRUE, stringsAsFactors = FALSE)
splinkData <- read.delim("speciesLink_all_49771_20190521013813.txt.out",
    header = TRUE, stringsAsFactors = FALSE)

### clean the SpeciesLink dataset remove data from
### specieslinkData with notes containing 'bloqueado'
splinkData <- splinkData[-grep(x = splinkData$notes, pattern = "bloqueado"),
    ]
# the latter case was messing with the type of the
# coordinate columns, coerce them to numeric
splinkData$longitude <- as.numeric(splinkData$longitude)
splinkData$latitude <- as.numeric(splinkData$latitude)
# coordinate automatic cleaning examine the boxplot
boxplot(splinkData$latitude)
boxplot(splinkData$longitude)
# filter out all coordinates east of (South
# America+Panama)'s easternmost point: -34.7930, João
# Pessoa, Paraíba, and
splinkData <- splinkData[-which(splinkData$longitude > -34.793),
    ]
```

```r
### Clean the GBIF dataset remove cases that are explicitly
### showing geospatial data
gbifData <- gbifData[-which(gbifData$hasGeospatialIssues ==
    "true"), ]
# remove all coordinates out of (South America + Panama)'s
# bounds
gbifData <- gbifData[-which(gbifData$decimalLongitude > -34.793),
    ]
gbifData <- gbifData[-which(gbifData$decimalLongitude < -83.0521),
    ]
gbifData <- gbifData[-which(gbifData$decimalLatitude > 12.4583),
    ]
# remove an erroneous record of Corydoras from Guajira
# along with caribbean marine occurrences
gbifData <- gbifData[-which(gbifData$decimalLatitude > 12.2),
    ]
# remove erroneous records from the Pacific
gbifData <- gbifData[-which(gbifData$decimalLatitude == 3.997866),
    ]
# remove erroneous records from the Caribbean
gbifData <- gbifData[-which(gbifData$decimalLatitude == 11.616667),
    ]
# remove erroneous records from the Caribbean
gbifData <- gbifData[-which(gbifData$decimalLatitude == 11),
    ]
# remove erroneous records from the Atlantic
gbifData <- gbifData[-which(gbifData$decimalLatitude == -16.851667),
    ]
# remove erroneous records from the Atlantic
gbifData <- gbifData[-which(gbifData$decimalLatitude == -24.134167),
    ]
# remove erroneous records from the Atlantic
gbifData <- gbifData[-which(gbifData$decimalLatitude == -23.983333),
    ]
# remove erroneous records from the Atlantic
gbifData <- gbifData[-which(gbifData$decimalLatitude == -24.134167),
    ]
# remove erroneous records from the Atlantic
gbifData <- gbifData[-which(gbifData$decimalLatitude == -29.943333),
    ]
# remove fossil occurrences
gbifData <- gbifData[-which(gbifData$basisOfRecord == "FOSSIL_SPECIMEN"),
    ]

### Clean the specieslink dataset remove an erroneous record
### of Corydoras from Guajira along with caribbean marine
### occurrences
splinkData <- splinkData[-which(splinkData$latitude > 12.2),
    ]
# remove erroneous records from the Atlantic
splinkData <- splinkData[-which(splinkData$latitude == -24.134167),
    ]
# remove erroneous records from the Atlantic
```

```r
splinkData <- splinkData[-which(splinkData$latitude == -29.943333),
    ]

### Taxonomic cleaning remove data w/o genus
gbifData <- gbifData[-which(gbifData$genus == ""), ]
splinkData <- splinkData[-which(splinkData$genus == ""), ]
splinkData <- splinkData[-which(splinkData$genus == "Callichthyidae"),
    ]

### subsetting construct the dataframe concatenating column
### contents and creating on the fly a column with
### information on the source of the record
callichCoords <- data.frame(genus = c(gbifData$genus, splinkData$genus),
    species = c(gbifData$species, splinkData$scientificname),
    latitude = c(gbifData$decimalLatitude, splinkData$latitude),
    longitude = c(gbifData$decimalLongitude, splinkData$longitude),
    source = c(rep("gbif", times = nrow(gbifData)), rep("splink",
        times = nrow(splinkData))), stringsAsFactors = FALSE)
# remove missing coordinates.
# identical(is.na(callichCoords$latitude),
# is.na(callichCoords$longitude)) evaluates to TRUE so we
# can just pick any of the coordinate components
callichCoords <- callichCoords[!is.na(callichCoords$longitude),
    ]
# write the clean dataset
write.table(x = callichCoords, file = "callichtyidae.tab",
    sep = "\t", row.names = FALSE, fileEncoding = "UTF-8")
```

##### S2.1.2 *Lepidosiren*

```r
gbifData <- read.delim("occurrence.txt", header = TRUE, stringsAsFactors = FALSE)
splinkData <- read.delim("speciesLink_all_49771_20190521013813.txt.out",
    header = TRUE, stringsAsFactors = FALSE)

### clean the SpeciesLink dataset coordinate automatic
### cleaning examine the boxplot
boxplot(splinkData$latitude)
boxplot(splinkData$longitude)
# filter out all coordinates east of (South
# America+Panama)'s easternmost point: -34.7930, João
# Pessoa, Paraíba, and
splinkData <- splinkData[-which(splinkData$longitude > -34.793),
    ]

### Clean the GBIF dataset remove cases that are explicitly
### showing geospatial data
gbifData <- gbifData[-which(gbifData$hasGeospatialIssues ==
    "true"), ]
# remove fossil occurrences
gbifData <- gbifData[-which(gbifData$basisOfRecord == "FOSSIL_SPECIMEN"),
    ]

### Taxonomic cleaning
```

```
### subsetting construct the dataframe concatenating column
### contents and creating on the fly a column with
### information on the source of the record
lepidoCoords <- data.frame(genus = c(gbifData$genus, splinkData$genus),
    species = c(gbifData$species, splinkData$scientificname),
    latitude = c(gbifData$decimalLatitude, splinkData$latitude),
    longitude = c(gbifData$decimalLongitude, splinkData$longitude),
    source = c(rep("gbif", times = nrow(gbifData)), rep("splink",
        times = nrow(splinkData))), stringsAsFactors = FALSE)
# remove missing coordinates.
# identical(is.na(lepidoCoords$latitude),
# is.na(lepidoCoords$longitude)) evaluates to TRUE so we
# can just pick any of the coordinate components
lepidoCoords <- lepidoCoords[!is.na(lepidoCoords$longitude),
    ]
# write the clean dataset
write.table(x = lepidoCoords, file = "lepidosiren.tab", sep = "\t",
    row.names = FALSE, fileEncoding = "UTF-8")
```

##### S2.1.3 *Phractocephalus*

```
gbifData <- read.delim("occurrence.txt", header = TRUE, stringsAsFactors = FALSE)
splinkData <- read.delim("speciesLink_all_49771_20190521013813.txt.out",
    header = TRUE, stringsAsFactors = FALSE)

### clean the SpeciesLink dataset examine the boxplot
boxplot(splinkData$latitude)
boxplot(splinkData$longitude)
# filter out all coordinates east of (South
# America+Panama)'s easternmost point: -34.7930, João
# Pessoa, Paraíba, and
splinkData <- splinkData[-which(splinkData$longitude > -34.793),
    ]

### Clean the GBIF dataset remove cases that are explicitly
### showing geospatial data
gbifData <- gbifData[-which(gbifData$hasGeospatialIssues ==
    "true"), ]
# remove all coordinates out of (South America + Panama)'s
# bounds
gbifData <- gbifData[-which(gbifData$decimalLongitude < -83.0521),
    ]
gbifData <- gbifData[-which(gbifData$decimalLatitude > 12.4583),
    ]
# remove fossil occurrences
gbifData <- gbifData[-which(gbifData$basisOfRecord == "FOSSIL_SPECIMEN"),
    ]
# two doubtful records removed, one b/c of geographic
# mismatch the other because of geographic uncertainty
# placing the record out of the known distribution
gbifData <- gbifData[-which(gbifData$occurrenceID == "086E2F34-26EC-4270-9CC8-771FFCCA737B"),
    ]
```

```
gbifData <- gbifData[-which(gbifData$occurrenceID == "BR:UEL:MZUEL-Peixes:18116"),
    ]
# the last case was duplicated in the specieslink dataset
splinkData <- splinkData[-which(splinkData$catalognumber ==
    "17189"), ]

### Taxonomic cleaning remove an erroneous record of
### Paracanthopoma
splinkData <- splinkData[-which(splinkData$genus == "Paracanthopoma"),
    ]
### subsetting construct the dataframe concatenating column
### contents and creating on the fly a column with
### information on the source of the record
phractoCoords <- data.frame(genus = c(gbifData$genus, splinkData$genus),
    species = c(gbifData$species, splinkData$scientificname),
    latitude = c(gbifData$decimalLatitude, splinkData$latitude),
    longitude = c(gbifData$decimalLongitude, splinkData$longitude),
    source = c(rep("gbif", times = nrow(gbifData)), rep("splink",
        times = nrow(splinkData))), stringsAsFactors = FALSE)
# remove missing coordinates.
# identical(is.na(phractoCoords$latitude),
# is.na(phractoCoords$longitude)) evaluates to TRUE so we
# can just pick any of the coordinate components
phractoCoords <- phractoCoords[!is.na(phractoCoords$longitude),
    ]
# write the clean dataset
write.table(x = phractoCoords, file = "phractocephalus.tab",
    sep = "\t", row.names = FALSE, fileEncoding = "UTF-8")
```

###### S2.1.4 Serrasalmidae

```
gbifData <- read.delim("occurrence.txt", header = TRUE, stringsAsFactors = FALSE)
splinkData <- read.delim("speciesLink_all_49771_20190521013813.txt.out",
    header = TRUE, stringsAsFactors = FALSE)

### clean the SpeciesLink dataset coordinate automatic
### cleaning examine the boxplot
boxplot(splinkData$latitude)
boxplot(splinkData$longitude)
# filter out all coordinates east of (South
# America+Panama)'s easternmost point: -34.7930, João
# Pessoa, Paraíba, and
splinkData <- splinkData[-which(splinkData$longitude > -34.793),
    ]

### Clean the GBIF dataset remove cases that are explicitly
### showing geospatial data
gbifData <- gbifData[-which(gbifData$hasGeospatialIssues ==
    "true"), ]
# remove all coordinates out of (South America + Panama)'s
# bounds
gbifData <- gbifData[-which(gbifData$decimalLongitude > -34.793),
```

```r
    ]
gbifData <- gbifData[-which(gbifData$decimalLongitude < -83.0521),
    ]
gbifData <- gbifData[-which(gbifData$decimalLatitude > 12.4583),
    ]
# remove serrasalmid occurrences in the ocean and west of
# the Andes in Peru where they are certainly erroneous
# these occurrences are said to be from Leticia, Colombia.
gbifData <- gbifData[-which(gbifData$decimalLatitude == 3.997866),
    ]
# remove an occurrence from 'Pibas' (most likely Pebas)
# that mapped west of the Andes
gbifData <- gbifData[-which(gbifData$decimalLatitude == -12.454263),
    ]
# remove an occurrence from west of the Andes of Piaractus
# from Sucre, introduced to the Magdalena-Cauca
gbifData <- gbifData[-which(gbifData$decimalLatitude == 9.166804),
    ]
# remove fossil occurrences
gbifData <- gbifData[-which(gbifData$basisOfRecord == "FOSSIL_SPECIMEN"),
    ]

### Taxonomic cleaning remove occurrences uncertain to family
### level
splinkData <- splinkData[-which(splinkData$genus == ""), ]
gbifData <- gbifData[-which(gbifData$genus == ""), ]
### subsetting construct the dataframe concatenating column
### contents and creating on the fly a column with
### information on the source of the record
serraCoords <- data.frame(genus = c(gbifData$genus, splinkData$genus),
    species = c(gbifData$species, splinkData$scientificname),
    latitude = c(gbifData$decimalLatitude, splinkData$latitude),
    longitude = c(gbifData$decimalLongitude, splinkData$longitude),
    source = c(rep("gbif", times = nrow(gbifData)), rep("splink",
        times = nrow(splinkData))), stringsAsFactors = FALSE)
# remove missing coordinates.
# identical(is.na(serraCoords$latitude),
# is.na(serraCoords$longitude)) evaluates to TRUE so we can
# just pick any of the coordinate components
serraCoords <- serraCoords[!is.na(serraCoords$longitude), ]
# write the clean dataset
write.table(x = serraCoords, file = "serrasalmidae.tab", sep = "\t",
    row.names = FALSE, fileEncoding = "UTF-8")
```

### S3 Correlates to community similarity

```r
### Recalculate the distance matrix for the reduced dataset
### as presented in the manuscript faunal similarity analyses
### for the Marakaipao fish fauna
dataset <- read.delim(file = "neogeneFishOccs.tab", stringsAsFactors = FALSE)

#################### only freswater taxa
```

```r
dataset <- dataset[which(dataset$Environment == "Freshwater" |
    dataset$Environment == "Both"), ]

### Solimões-Pebas does not seem to be a fauna but a
### collection of different faunas across the Amazon, remove
### it
dataset <- dataset[, -grep(pattern = "Pebas", x = colnames(dataset))]

comMatrix <- dataset[, -c(1, 3, 18)]
rownames(comMatrix) <- comMatrix$Taxon
comMatrix <- comMatrix[-1]
comMatrix <- t(comMatrix)

### number of occurrences per fauna
sort(x = apply(X = comMatrix, MARGIN = 1, FUN = sum), decreasing = TRUE)
```

```
##           Urumaco            La.Venta           Rio.Acre          Ituzaingo
##                13                  11                  8                  6
##        Makaraipao         Fitzcarrald          Contamana           Castillo
##                 4                   4                  4                  3
##     Loyola.Mangan            Rio.Yuca           Utuquina Castilletes.marine
##                 2                   2                  1                  0
##           Pirabas            Cantaure
##                 0                   0
```

```r
### include only those faunas with at least the number of
### occurrences of Makaraipao
selectFaunas <- names(which(apply(X = comMatrix, MARGIN = 1,
    FUN = sum) >= apply(X = comMatrix, MARGIN = 1, FUN = sum)["Makaraipao"]))
comMatrix <- comMatrix[selectFaunas, ]

### remove zero-sum species after faunal selection
selectSpp <- names(which(apply(X = comMatrix, MARGIN = 2, FUN = sum) >
    0))
comMatrix <- comMatrix[, selectSpp]

### calculate the distance matrix using Bray-Curtis' method
distMatrixBray <- vegan::vegdist(comMatrix, method = "bray",
    binary = TRUE)
# rename labels in order to replace dots with spaces
attr(distMatrixBray, "Labels") <- gsub(pattern = "\\.", replacement = " ",
    x = attr(distMatrixBray, "Labels"))
# rename trans-Andean labels in order to place a leading
# asterisk
attr(distMatrixBray, "Labels") <- gsub(pattern = "Urumaco",
    replacement = "( T ) Urumaco", x = attr(distMatrixBray,
        "Labels"), fixed = TRUE)
attr(distMatrixBray, "Labels") <- gsub(pattern = "La Venta",
    replacement = "( T ) La Venta", x = attr(distMatrixBray,
        "Labels"), fixed = TRUE)
attr(distMatrixBray, "Labels") <- gsub(pattern = "Makaraipao",
    replacement = "( T ) Makaraipao", x = attr(distMatrixBray,
        "Labels"), fixed = TRUE)
```

```r
### similarity vs. linear distance
points <- sf::st_read("paleoMiocene.kml")
```

```
## Reading layer `paleoMio' from data source
##   `/home/balleng/Dropbox/Gustavo/papers/makaraipao/datasets/paleoMiocene.kml'
##   using driver `KML'
## Simple feature collection with 7 features and 2 fields
## Geometry type: POINT
## Dimension:     XYZ
## Bounding box:  xmin: -75.16058 ymin: -31.88504 xmax: -60.32894 ymax: 11.90889
## z_range:       zmin: 0 zmax: 0
## Geodetic CRS:  WGS 84
```

```r
# solve problems with labels
pairwisePoints <- sf::st_distance(points)
```

```
## st_as_s2(): dropping Z and/or M coordinate
## st_as_s2(): dropping Z and/or M coordinate
```

```r
rownames(pairwisePoints) <- points$Name
rownames(pairwisePoints) <- gsub(pattern = " ", replacement = ".",
    x = rownames(pairwisePoints))
rownames(pairwisePoints) <- gsub(pattern = ".Fm.", replacement = "",
    x = rownames(pairwisePoints))
rownames(pairwisePoints) <- gsub(pattern = "Acre", replacement = "Rio.Acre",
    x = rownames(pairwisePoints))
colnames(pairwisePoints) <- points$Name
colnames(pairwisePoints) <- gsub(pattern = " ", replacement = ".",
    x = colnames(pairwisePoints))
colnames(pairwisePoints) <- gsub(pattern = ".Fm.", replacement = "",
    x = colnames(pairwisePoints))
pairwisePoints <- as.dist(pairwisePoints)

# sort labels
pairwisePoints <- as.matrix(pairwisePoints)
pairwisePoints <- as.dist(pairwisePoints[order(rownames(pairwisePoints)),
    order(colnames(pairwisePoints))])

# recalculate the pairwise community distance for easy
# manipulation
pairwiseComm <- dist(comMatrix, method = "binary")

# sort labels
pairwiseComm <- as.matrix(pairwiseComm)
pairwiseComm <- as.dist(pairwiseComm[order(rownames(pairwiseComm)),
    order(colnames(pairwiseComm))])

# spearman's rho correlation
corPointsComm <- cor.test(x = pairwisePoints, y = pairwiseComm,
    method = "spearman", exact = FALSE)

plot(x = pairwisePoints, y = pairwiseComm, xlab = "Geographic distance",
    ylab = "Community similarity", xlim = c(0, 3500000), pch = 21,
    bg = "black")
```

```r
legend(x = "bottomright", legend = paste("rho = ", round(corPointsComm$estimate,
    digits = 3), "; p = ", round(corPointsComm$p.value, digits = 3),
    sep = ""))
```

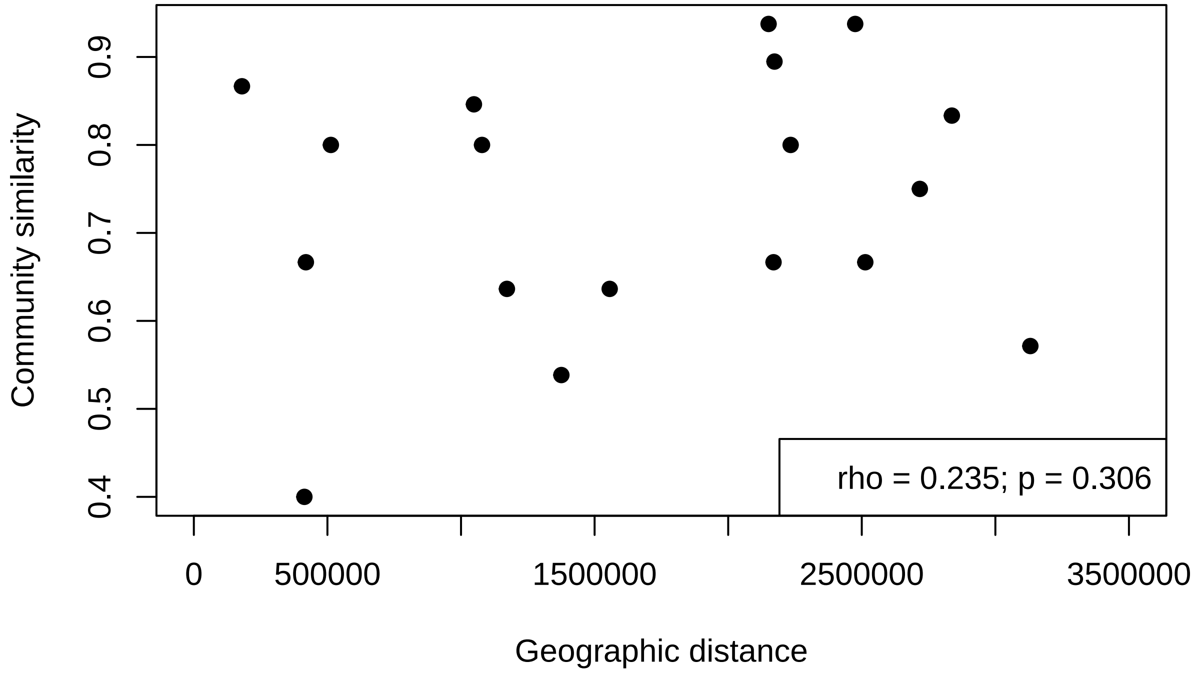

```r
# similaraity vs. geoAge

# age data Urumaco (Urumaco Fm.) ~ 8Ma; 8.0 Makaraipao
# (Castilletes Fm.) ~ 15 Ma; 15.0 La Venta (Honda Gr.) ~ 11
# - 13 Ma; 12.0 Contamana (Pebas-Ipururo Fms.) = middle to
# late Miocene ~ 14-8 Ma?; 11.0 Fitzcarrald (No
# stratigraphic control, Pebas Fm.?) = Laventan ~ 11.8 -
# 13.8 Ma; 12.8 Ma Ituzaingo (Conglomerado Osífero,
# Ituzaingo Fm.) ~ 6 - 9 Ma; 7.5 Rio Acre (Solimões Fm.) =
# Huayquerian ~ 6.8 - 9 Ma; 7.9

# Sorted: 07.5 Ituzaingo; 07.9 Rio Acre; 08.0 Urumaco; 11.0
# Contamana; 12.0 La Venta; 12.8 Fitzcarrald; 15.0
# Makaraipao;

geoAges <- c(11, 12.8, 7.5, 12, 15, 7.9, 8)

names(geoAges) <- c("Contamana", "Fitzcarrald", "Ituzaingo",
    "La.Venta", "Makaraipao", "Rio.Acre", "Urumaco")

pairwiseAges <- dist(geoAges)

# spearman's rho correlation
corAgesComm <- cor.test(x = pairwiseAges, y = pairwiseComm,
    method = "spearman", exact = FALSE)

plot(x = pairwiseAges, y = pairwiseComm, xlab = "Geological age distance",
```

```r
    ylab = "Community similarity", pch = 21, bg = "black")

legend(x = "bottomright", legend = paste("rho = ", round(corAgesComm$estimate,
    digits = 3), "; p = ", round(corAgesComm$p.value, digits = 3),
    sep = ""))
```

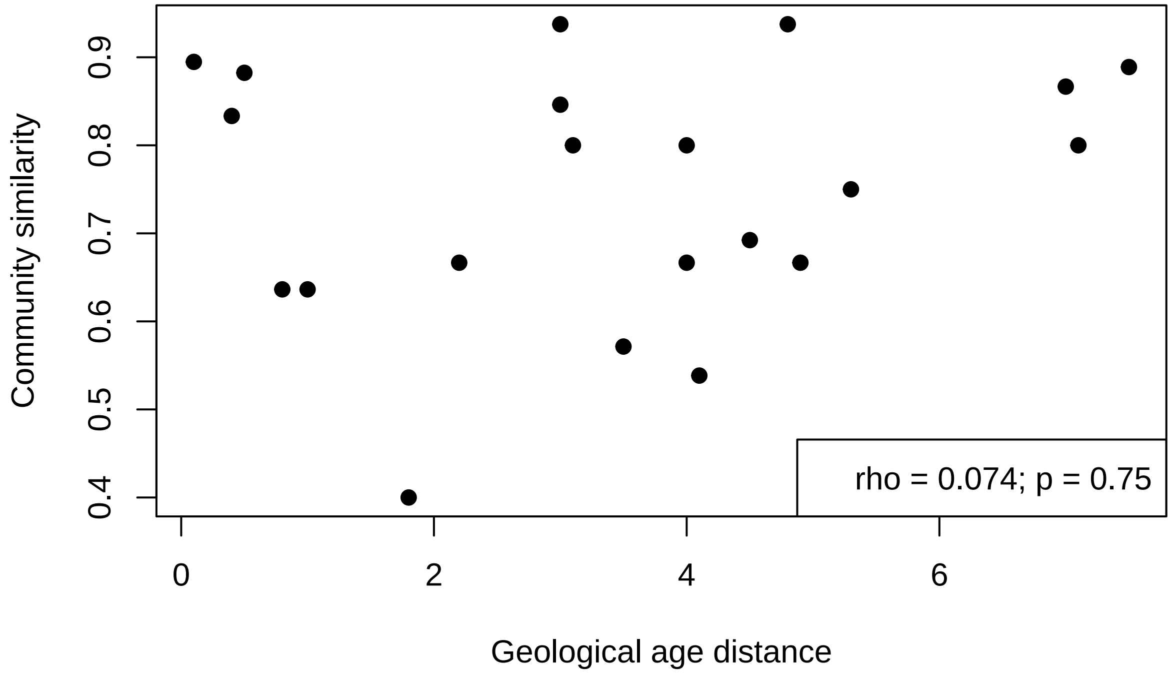

#### S4 Raw occurrence data

Table S1: Occurrence table. Acronyms of fossil faunas are: Castillo=CS, La Venta=LV, Makaraipao=MK, Castilletes marine=CM, Loyola-Mangan=LM, Rio Acre=RA, Solimões-Pebas=SP, Urumaco=UF, Utuquina=UQ, Pirabas=PI, Cantaure=CA, Ituzaingo=IT, Rio Yuca=RY, Fitzcarrald=FZ, Contamana=CT

| Family | Taxon | Environment | CS | LV | MK | CM | LM | RA | SP | UF | UQ | PI | CA | IT | RY | FZ | CT | Ref. |
|---|---|---|---|---|---|---|---|---|---|---|---|---|---|---|---|---|---|---|
| Acregoliathidae | Acregoliath | Freshwater | 0 | 1 | 0 | 0 | 0 | 1 | 0 | 0 | 0 | 0 | 0 | 0 | 0 | 1 | 0 | Ballen and Moreno-Bernal (2019); Lundberg et al. (2010); Tejada-Lara et al. (2015) |
| Arapaimidae | Arapaima | Freshwater | 0 | 1 | 0 | 0 | 0 | 1 | 0 | 0 | 0 | 0 | 0 | 0 | 0 | 0 | 0 | Lundberg et al. (2010) |
| Anostomidae | Leporinus | Freshwater | 0 | 1 | 0 | 0 | 1 | 0 | 1 | 0 | 0 | 0 | 0 | 1 | 0 | 0 | 1 | Antoine et al. (2016); Bogan et al. (2012); Lundberg et al. (2010) |
| Characidae | Salminus | Freshwater | 0 | 0 | 0 | 0 | 0 | 0 | 0 | 0 | 0 | 0 | 0 | 1 | 0 | 0 | 0 | Cione and Azpelicueta (2013) |
| Serrasalmidae | Colossoma | Freshwater | 1 | 1 | 0 | 0 | 0 | 1 | 0 | 1 | 0 | 0 | 0 | 1 | 0 | 0 | 0 | Cione et al. (2000); Lundberg et al. (2010) |
| Serrasalmidae | Megapiranha | Freshwater | 0 | 0 | 0 | 0 | 0 | 0 | 0 | 0 | 0 | 0 | 0 | 1 | 0 | 0 | 0 | Cione et al. (2009) |
| Serrasalmidae | Mylossoma | Freshwater | 1 | 0 | 1 | 0 | 0 | 0 | 0 | 0 | 0 | 0 | 0 | 0 | 0 | 0 | 0 | Lundberg et al. (2010) |
| Serrasalmidae | Piaractus | Freshwater | 0 | 0 | 1 | 0 | 0 | 0 | 0 | 1 | 0 | 0 | 0 | 0 | 0 | 0 | 0 | Lundberg et al. (2010) |
| Cynodontidae | Hydrolycus | Freshwater | 0 | 1 | 0 | 0 | 0 | 0 | 0 | 0 | 0 | 0 | 0 | 1 | 0 | 1 | 1 | Antoine et al. (2016); Cione et al. (2000); Lundberg et al. (2010); Tejada-Lara et al. (2015) |
| Erythrinidae | Paleohoplias | Freshwater | 0 | 0 | 0 | 0 | 0 | 1 | 0 | 0 | 0 | 0 | 0 | 0 | 0 | 0 | 0 | Lundberg et al. (2010) |
| Erythrinidae | Hoplias | Freshwater | 0 | 1 | 0 | 0 | 1 | 0 | 1 | 0 | 0 | 0 | 0 | 0 | 0 | 0 | 0 | Lundberg et al. (2010) |
| Callichthyidae | Hoplosternum | Freshwater | 0 | 1 | 0 | 0 | 0 | 1 | 0 | 0 | 0 | 0 | 0 | 0 | 0 | 0 | 0 | Lundberg et al. (2010) |
| Loricariidae | Acanthicus | Freshwater | 0 | 1 | 0 | 0 | 0 | 0 | 0 | 1 | 0 | 0 | 0 | 0 | 0 | 0 | 0 | Lundberg et al. (2010) |
| Doradidae | Doraops | Freshwater | 0 | 0 | 0 | 0 | 0 | 0 | 0 | 1 | 0 | 0 | 0 | 0 | 0 | 0 | 0 | Lundberg et al. (2010) |
| Doradidae | Doras | Freshwater | 0 | 0 | 0 | 0 | 0 | 0 | 0 | 1 | 0 | 0 | 0 | 0 | 0 | 0 | 0 | Lundberg et al. (2010) |
| Doradidae | Oxydoras | Freshwater | 0 | 0 | 0 | 0 | 0 | 0 | 0 | 0 | 1 | 0 | 0 | 0 | 0 | 0 | 0 | Lundberg et al. (2010) |
| Doradidae | Rhinodoras | Freshwater | 0 | 0 | 0 | 0 | 0 | 0 | 0 | 1 | 0 | 0 | 0 | 0 | 0 | 0 | 0 | Lundberg et al. (2010) |
| Pimelodidae | Brachyplatystoma | Freshwater | 0 | 1 | 0 | 0 | 0 | 0 | 0 | 1 | 0 | 0 | 0 | 0 | 0 | 0 | 0 | Aguilera et al. (2013a); Lundberg et al. (2010) |
| Pimelodidae | Phractocephalus | Freshwater | 0 | 1 | 1 | 0 | 0 | 1 | 0 | 1 | 0 | 0 | 0 | 1 | 1 | 1 | 1 | Antoine et al. (2016); Azpelicueta and Cione (2016); Lundberg et al. (2010); Tejada-Lara et al. (2015) |
| Pimelodidae | Platysilurus | Freshwater | 0 | 0 | 0 | 0 | 0 | 0 | 0 | 1 | 0 | 0 | 0 | 0 | 1 | 0 | 0 | Lundberg et al. (2010) |
| Pimelodidae | Zungaro | Freshwater | 0 | 0 | 0 | 0 | 0 | 1 | 0 | 0 | 0 | 0 | 0 | 0 | 0 | 0 | 0 | Lundberg et al. (2010) |
| Ariidae | Amphiarius | Marine | 0 | 0 | 0 | 0 | 0 | 0 | 0 | 1 | 0 | 0 | 0 | 0 | 0 | 0 | 0 | Lundberg et al. (2010) |
| Ariidae | Aspistor | Both | 0 | 0 | 0 | 0 | 0 | 0 | 0 | 1 | 0 | 0 | 0 | 0 | 0 | 0 | 0 | Lundberg et al. (2010) |
| Ariidae | Bagre | Marine | 1 | 0 | 0 | 1 | 0 | 0 | 0 | 1 | 0 | 0 | 1 | 0 | 0 | 0 | 0 | Aguilera et al. (2013b); Lundberg et al. (2010) |
| Ariidae | Cathorops | Marine | 0 | 0 | 0 | 0 | 0 | 0 | 0 | 0 | 0 | 1 | 0 | 0 | 0 | 0 | 0 | Aguilera et al. (2013b) |
| Ariidae | Cantarius | Marine | 0 | 0 | 0 | 1 | 0 | 0 | 0 | 0 | 0 | 0 | 1 | 0 | 0 | 0 | 0 | Aguilera et al. (2013b) |
| Ariidae | Notarius | Marine | 0 | 0 | 0 | 0 | 0 | 0 | 0 | 1 | 0 | 0 | 0 | 0 | 0 | 0 | 0 | Lundberg et al. (2010) |
| Ariidae | Sciades | Both | 0 | 0 | 0 | 0 | 0 | 0 | 0 | 1 | 0 | 0 | 0 | 0 | 0 | 0 | 0 | Lundberg et al. (2010) |
| Sciaenidae | Ctenosciaena | Marine | 1 | 0 | 0 | 0 | 0 | 0 | 0 | 0 | 0 | 0 | 0 | 0 | 0 | 0 | 0 | Lundberg et al. (2010) |
| Sciaenidae | Cynoscion | Both | 0 | 0 | 0 | 0 | 0 | 0 | 0 | 1 | 0 | 0 | 0 | 0 | 0 | 0 | 0 | Lundberg et al. (2010) |
| Sciaenidae | Equetus | Marine | 1 | 0 | 0 | 0 | 0 | 0 | 0 | 1 | 0 | 0 | 0 | 0 | 0 | 0 | 0 | Lundberg et al. (2010) |
| Sciaenidae | Larimus | Marine | 0 | 0 | 0 | 0 | 0 | 0 | 0 | 1 | 0 | 0 | 0 | 0 | 0 | 0 | 0 | Lundberg et al. (2010) |
| Sciaenidae | Micropogonias | Marine | 0 | 0 | 0 | 0 | 0 | 0 | 0 | 1 | 0 | 0 | 0 | 0 | 0 | 0 | 0 | Lundberg et al. (2010) |
| Sciaenidae | Nebris | Marine | 0 | 0 | 0 | 0 | 0 | 0 | 0 | 1 | 0 | 0 | 0 | 0 | 0 | 0 | 0 | Lundberg et al. (2010) |
| Sciaenidae | Ophioscion | Marine | 0 | 0 | 0 | 0 | 0 | 0 | 0 | 1 | 0 | 0 | 0 | 0 | 0 | 0 | 0 | Lundberg et al. (2010) |
| Sciaenidae | Pachypops | Freshwater | 0 | 0 | 0 | 0 | 0 | 0 | 1 | 0 | 0 | 0 | 0 | 0 | 0 | 0 | 0 | Lundberg et al. (2010) |
| Sciaenidae | Paralonchurus | Marine | 1 | 0 | 0 | 0 | 0 | 0 | 0 | 0 | 0 | 0 | 0 | 0 | 0 | 0 | 0 | Lundberg et al. (2010) |
| Sciaenidae | Plagioscion | Freshwater | 1 | 0 | 0 | 0 | 0 | 0 | 1 | 1 | 0 | 0 | 0 | 0 | 0 | 0 | 0 | Lundberg et al. (2010) |
| Sciaenidae | Protosciaena | Marine | 1 | 0 | 0 | 0 | 0 | 0 | 0 | 0 | 0 | 0 | 0 | 0 | 0 | 0 | 0 | Lundberg et al. (2010) |
| Sciaenidae | Xenotolithus | NA | 0 | 0 | 0 | 0 | 0 | 0 | 1 | 0 | 0 | 0 | 0 | 0 | 0 | 0 | 0 | Lundberg et al. (2010) |
| Serranidae | Epinephelus | Marine | 0 | 0 | 0 | 0 | 0 | 0 | 0 | 1 | 0 | 0 | 0 | 0 | 0 | 0 | 0 | Lundberg et al. (2010) |
| Sphyraenidae | Sphyraena | Marine | 1 | 0 | 0 | 0 | 0 | 0 | 0 | 0 | 0 | 0 | 0 | 0 | 0 | 0 | 0 | Lundberg et al. (2010) |
| Scombridae | Acanthocybium | Marine | 1 | 0 | 0 | 0 | 0 | 0 | 0 | 0 | 0 | 0 | 0 | 0 | 0 | 0 | 0 | Lundberg et al. (2010) |
| Lepidosirenidae | Lepidosiren | Freshwater | 0 | 1 | 1 | 0 | 0 | 1 | 0 | 0 | 0 | 0 | 0 | 0 | 0 | 1 | 1 | Antoine et al. (2016); Lundberg et al. (2010); Tejada-Lara et al. (2015) |

### References

Aguilera, O. A., Lundberg, J. G., Birindelli, J. L. O., Sabaj Pérez, M. H., Jaramillo, C. A., and Sánchez-Villagra, M. R. (2013a). Palaeontological evidence for the last temporal occurrence of the ancient Western Amazonian river outflow into the Caribbean. *PLoS ONE*, 8(9):1–17.

Aguilera, O. A., Moraes-Santos, H., Costa, S. A. R. F., Ohe, F., Jaramillo, C. A., and Nogueira, A. (2013b). Ariid sea catfishes from the coeval Pirabas (Northeastern Brazil), Cantaure, Castillo (Northwestern Venezuela), and Castilletes (North Colombia) formations (early Miocene), with description of three new species. *Swiss Journal of Palaeontology*, 132(1):45–68.

Antoine, P.-O., Abello, M. A., Adnet, S., Altamirano Sierra, A. J., Baby, P., Billet, G., Boivin, M., Calderón, Y., Candela, A., Chabain, J., Corfu, F., Croft, D. A., Ganerød, M., Jaramillo, C. A., Klaus, S., Marivaux, L., Navarrete, R. E., Orliac, M. J., Parra, F., Pérez, M. E., Pujos, F., Rage, J. C., Ravel, A., Robinet, C., Roddaz, M., Tejada-Lara, J. V., Vélez-Juarbe, J., Wesselingh, F. P., and Salas-Gismondi, R. (2016). A 60-million-year Cenozoic history of western Amazonian ecosystems in Contamana, eastern Peru. *Gondwana Research*, 31:30–59.

Azpelicueta, M. d. l. M. and Cione, A. L. (2016). A southern species of the tropical catfish genus *Phractocephalus* (Teleostei: Siluriformes) in the Miocene of South America. *Journal of South American Earth Sciences*, 67:221–230.

Ballen, G. A. and Moreno-Bernal, J. W. (2019). New records of the enigmatic Neotropical fossil fish *Acregoliath rancii* (Teleostei *incertae sedis*) from the middle Miocene Honda group of Colombia. *Ameghiniana*, 56(6):431–440.

Bogan, S., Sidlauskas, B., Vari, R. P., and Agnolin, F. L. (2012). *Arrhinolemur scalabrinii* Ameghino, 1898, of the late Miocene - a taxonomic journey from the Mammalia to the Anostomidae (Ostariophysi: Characiformes). *Neotropical Ichthyology*, 10(3):555–560.

Cione, A. L. and Azpelicueta, M. d. l. M. (2013). The first fossil species of *Salminus*, a conspicuous South American freshwater predatory fish (Teleostei , Characiformes), found in the Miocene of Argentina. *Journal of Vertebrate Paleontology*, 33(5):1051–1060.

Cione, A. L., Azpelicueta, M. d. l. M., Bond, M., Carlini, A. A., Casciotta, J. R., Cozzuol, M. A., de la Fuente, M., Gasparini, Z., Goin, F. J., Noriega, J., Scillato-Yané, G. J., Soibelzon, L., Tonni, E. P., Verzi, D., and Vucetich, M. G. (2000). The Miocene vertebrates from Paraná, eastern Argentina Miocene vertebrates from Entre Ríos province, eastern Argentina. *Correlación Geológica*, 14:191–237.

Cione, A. L., Dahdul, W. M., Lundberg, J. G., and Machado-Allison, A. (2009). *Megapiranha paranensis*, a new genus and species of Serrasalmidae (Characiformes, Teleostei) from the upper Miocene of Argentina. *Journal of Vertebrate Paleontology*, 29(2):350–358.

Lundberg, J. G., Sabaj Pérez, M. H., Dahdul, W. M., and Aguilera, O. A. (2010). The Amazonian Neogene fish fauna. In Hoorn, C. and Wesselingh, F. P., editors, *Amazonia, Landscape and Species Evolution: A Look Into the Past*, pages 281–301. Blackwell Publishing.

Tejada-Lara, J. V., Salas-Gismondi, R., Pujos, F., Baby, P., Benammi, M., Brusset, S., de Franceschi, D., Espurt, N., Urbina, M., and Antoine, P.-O. (2015). Life in proto-Amazonia: Middle Miocene Mammals from the Fitzcarrald Arch (Peruvian Amazonia). *Palaeontology*, 58:341–378.
